# A comprehensive benchmarking study of protein structure alignment tools based on downstream task performance

**DOI:** 10.1101/2025.03.11.642719

**Authors:** Zhuoyang Chen, Xuechen Zhang, Weichuan Yu, Qiong Luo

## Abstract

In this study, we investigate the performance of nine protein structure alignment tools by analyzing the influence of alignment results in accomplishing three downstream biological tasks: homology detection, phylogeny reconstruction, and function inference. These tools include (1) traditional sequential methods using both 3D and 2D structure representations, (2) non-sequential methods, (3) flexible methods, and (4) deep-learning methods. Canonical sequence alignment methods Needleman-Wunsch algorithm and BLASTp are used as baseline methods. We show that accuracies in downstream tasks can be uncorrelated with alignment quality metrics such as TM-score and RMSD, highlighting the discrepancy between the alignment results and the purposes of using them. We identify scenarios where structure alignment results outperform sequence alignment results. In homology detection, structure-based methods are much better than sequence alignment. In phylogeny reconstruction, structure-based methods generally outperform sequence-based methods on the filtered dataset with proteins sharing low sequence similarity. Moreover, we show that structure information improves the overall performance of these tools when used together with sequence information in phylogeny reconstruction and function inference. We also test the running time and CPU/GPU memory consumption of these tools for a large number of queries. Our study suggests that biological problems that were previously addressed with sequence-based methods using only sequence information could be further improved by using structure information alone or using both sequence and structure information. The trade-off between task accuracy and speed is the major consideration in developing new alignment tools for downstream tasks. We recommend both TMalign and KPAX for these tasks because of their good balance between running time and memory consumption, and relatively good and stable accuracy performance in downstream tasks. In tasks that require a large number of pairwise comparisons, such as homology detection and function inference, traditional methods outperform DL methods at the cost of long running time, and Foldseek is the best choice to achieve relatively high accuracy in a reasonable time.

**Graphical abstract:** Fig. 1:
Graphical abstract.

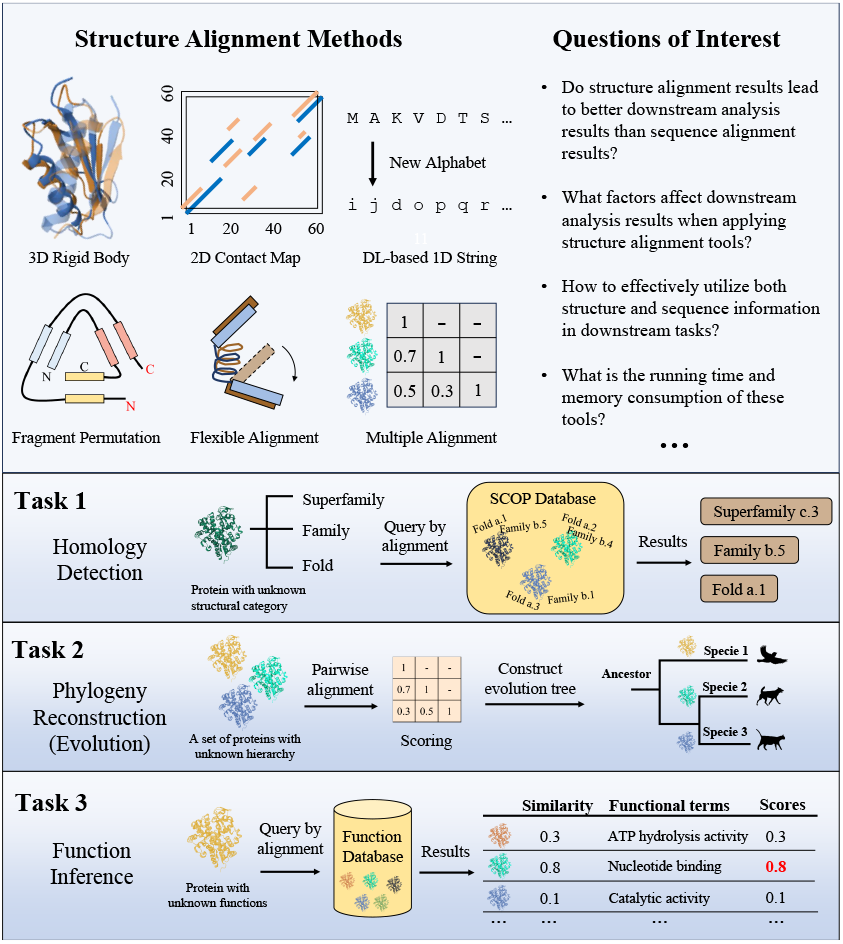

## 2 Introduction

Protein structure alignment is a fundamental problem in biology. It is a critical step in many tasks, including drug/enzyme design, homology detection (structural classification), protein structure modeling, and protein function annotation (1). Despite the increasing number of resolved protein structures documented in the PDB database (2), most protein-related tasks, such as phylogeny (evolutionary hierarchy) reconstruction and function prediction, were previously performed using only amino acid sequence information only. We name these tasks as sequence-based tasks. Although protein structure is commonly believed to be more conserved and provides more information than sequence in terms of evolution (3) and function (4), few studies have investigated whether using structure information alone can outperform using sequence information, or provided insight into how to use both sequence and structure information for better performance (5, 6) in these sequence-based tasks. In addition, few studies have comprehensively assessed the accuracies of various downstream biological tasks such as phylogeny reconstruction and function inference using existing structure alignment tools. Most recent tools focus only on the homology detection task that infers the structural category of a protein based on its structural overlap with proteins that have annotated category labels in the database. Researchers have noticed that tools that produce the greatest structural overlap or the most accurate structure-based alignment patterns do not necessarily yield the highest accuracy in downstream tasks (7, 8, 9). How such discrepancies are reflected in various structure alignment tools and what factors contribute most are yet to be explored, especially for real-life biological applications. This lack of investigation hinders people’s awareness of the importance of developing protein structure alignment tools and incorporating protein structure information as a crucial factor in developing biomedicine targeting models.

Two main challenges prohibit the exploration of utilizing structure information in sequence-based tasks: the lack of full-length structures from sequences and the lack of fast structure alignment tools for large-scale queries (millions of comparisons). Protein structures can be resolved by X-ray crystallography, cryo-electron microscopy, nuclear magnetic resonance (NMR) spectroscopy, and other methods (2). Such processes are time-consuming and costly, and usually only produce fragments of data instead of the entire protein structure. As structure prediction methods such as AlphaFold (10) generate millions of publicly available protein structures, querying these databases becomes the performance bottleneck. Various protein structure alignment tools have been developed in the last two decades, but these traditional tools are too slow for a large number of queries. It is reported that querying a single structure takes months to traverse the AlphaFold database AlphaFoldDB, the largest protein structure database containing over 200 million structures (11). Such a long query time is prohibitive for the evaluation of real-world applications. Only recently have several deep learning (DL)-based structure alignment methods been designed for large-scale queries (12, 7). With millions of AlphaFold-predicted full-length protein structures and fast structure search tools, we aim to conduct a comprehensive performance comparison on protein structure alignment tools that employ various technical schemes including (1) traditional methods (DALI (13), TMalign (14), and

DeepAlign (15)), (2) a non-sequential method USalign2 (16) that allows motif/fragment permutations, (3) a flexible method KPAX (17) that utilizes spatially local features, and (4) DL methods (DeepBLAST (7), pLM-BLAST (18), and Foldseek (12)) across three tasks. We also included multiple alignment tools (Clustal Omega (19), mTMalign (20), 3DCOMB (21), KPAX-flex (22), and Foldmason (23)) when reconstructing the phylogenetic trees.

Traditional sequence alignment methods including Needleman-Wunsch (24) and BLASTp (25) are used as baselines. A summary of the methods and tools included is provided in Table 1.

**Table 1.**
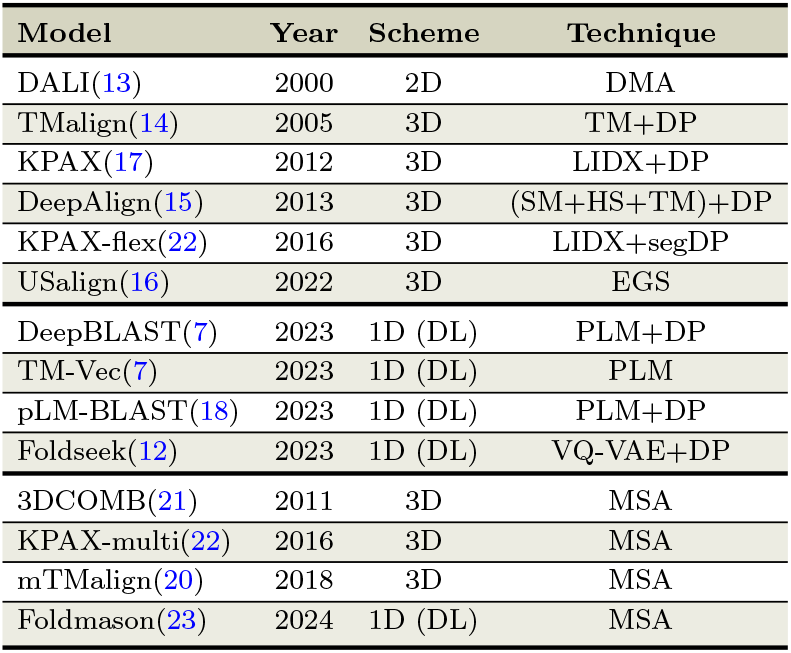
List of protein structure alignment tools included in this study. **DMA:** Distance Matrix Alignment. **TM:** TM-score. **DP:** Dynamic Programming. **LIDX:** Local Indexing. **SM:** Substitution Matrix. **HB:** Hydrogen-binding Score. **segDP:** Segmented Dynamic Programming. **EGS:** Enhanced Greedy Search. **DL:** Deep Learning. **PLM:** Protein Language Model. **VQ-VAE:** Vector-Quantized Variational AutoEncoder. **MSA:** Multiple Structure Alignment.

Homology detection, phylogeny reconstruction, and protein function inference reflect the fundamental question in biology, namely the relationship between protein structure, function, and evolution. The relationship between protein function and structure is much closer than that between function and sequence (4). Similarly, the relationship between phylogeny (evolutionary hierarchy) and protein structure is closer than that between phylogeny and sequence across species (3). In these tasks, finding exact alignment patterns is no longer the primary goal, but rather the relative similarity among a set of proteins. Although most structure alignment tools mainly focus on deriving the largest structural overlap between two structures, the alignment task itself is only an intermediate step for many downstream tasks in the biomedicine field. For example, structure alignment can be used to search for a similar protein based on structure for novel enzyme design and structure prediction, but the design and prediction performance could be less correlated with the structure alignment accuracy. Thus, as more and more full-length protein structures and advanced structure alignment tools are available, it is necessary to investigate how to effectively utilize both structure and sequence information to improve the downstream task accuracy when applying structure alignment methods.

On utilizing structure information, Carpentier et. al. find that multiple sequence alignment (MSA) patterns generated by structure-based methods are more accurate than those generated by sequence-based and sequence plus structure methods when sequence identities (sequence similarity) are low in a certain protein family (5). The authors suggest that a model combining sequence and structure evolutionary information could achieve better results in downstream tasks, and such a model is difficult to design. Some studies evaluate the performance of applying structure alignment results in homology detection but these studies do not include a comprehensive spectrum of methods and tasks (7, 18). Balaji et. al. show that evolutionary trees reconstructed from sequence and structure can be very different, and structure-based trees can be more accurate than sequence-based trees especially when sequence identities are low (26). However, their study does not focus on evaluating the accuracy of trees, but mainly on topology differences instead, and the evaluation is based on a structural dataset but not consider much evolution information. The Foldtree study (6) on phylogeny reconstruction demonstrates the potential of combining sequence and structure information for phylogeny inference using Foldseek, but it does not focus on comprehensively comparing different alignment methods. Few benchmark studies on functional inference exist for structure alignment methods due to the lack of full-length structure data and long running time for large-scale queries with traditional non-DL tools. Only one recent tool GOCurator, the top-1 solution of CAFA5, incorporates Foldseek as a component, and the authors of GOCurator found that BLASTp outperforms Foldseek as the homology search strategy. Few studies include non-sequential or flexible methods for comparison or use full-length structure data.

In this study, we conduct a benchmarking study including more downstream tasks and a greater diversity of alignment tools. First, we extend the downstream tasks of structure alignment methods from homology detection to phylogeny reconstruction and function inference, given the availability of vast predicted full-length protein structures in AlphaFoldDB (11). Second, we include structure alignment tools in a wide spectrum of schemes, including sequential/non-sequential, flexible, and DL-based methods, and also use traditional sequence alignment methods as baselines. Through such a comprehensive benchmarking study, we explore the utilization of structure data in tasks that previously relied on sequence homology search. Specifically, we find out in what situations structure alignment outperforms sequence alignment in these tasks and what factors contribute to the performance advantage. For each task, we provide suggestions for alignment tool selection and share important considerations for performance improvement.

The contributions of our work include the following:

1. This is the first benchmarking study on protein structure alignment methods for various downstream tasks.
2. We conduct a comprehensive comparison that includes a wide spectrum of tools under various schemes, such as pairwise/multiple alignment, rigid/flexible, sequential/non-sequential, and traditional/DL-based methods.
3. We demonstrate the importance of utilizing both sequence and structure information for better performance in downstream tasks and show that such a combination can be applied to traditional structure alignment tools.
4. We investigate factors that affect the performance of various tools and find that the type of scoring metrics, the mechanism for partial alignment, and the quality of AlphaFold predicted structures are great contributors.

## 3 Materials and methods

### 3.1 Datasets

#### 3.1.1 Malidup

Malidup (27) includes a group of proteins that have internal duplication within the same chain. It consists of 241 pairwise structure alignments for homologous domains. About one half of these pairs are remote homologs. Alignment results of TMalign, DALI, and manual inspection have been provided in the dataset.

#### 3.1.2 Malisam

Malisam (28) consists of 130 pairs of analogous motifs and each pair of proteins are structural analogs. More than 60% of these pairs belong to different folds as defined in SCOP. This dataset is more challenging than Malidup, as the sequence identity of any two proteins is very low and the two proteins in a pair could be non-homologous. Alignment results of TMalign, DALI, and manual inspection have been provided in the dataset.

#### 3.1.3 SCOP140

SCOP140 is constructed based on the SCOPe 2.07 database (29) and is used by DaliLite v.5 (13) to benchmark fold detection across different protein structure alignment tools. It contains 140 protein structures as the test set and 15211 structures as the target set. Each protein has a category label, annotating fold, family, and superfamily information. A pair of proteins with one from the test set (query protein) and the other from the target set (target protein) are aligned and assessed to predict whether they share the same category label. Accuracy results of DALI, TMalign, and DeepAlign have been provided in the dataset.

#### 3.1.4 SwissTree

SwissTree (30) is a database containing 19 manually annotated phylogenic trees, each constituting a protein family. We only use the first 11 trees in the database according to the Foldtree settings (6).

#### 3.1.5 CAFA3-MF

The Critical Assessment of Functional Annotation (CAFA) is a competition on predicting the functions of unannotated proteins (31). Based on the Gene Ontology (GO) definition (32), protein functions can be categorized by three GO aspects: Biological Process (BP), Molecular Function (MF), and Cellular Component (CC). Data is downloaded from the TEMPROT paper (33) with the prefix *mf*. We choose the MF aspect because it has the least but sufficient number of query and target proteins, which reduces the running time for our benchmarking study. The CAFA3-MF query set contains 1137 proteins and the target set contains 32421 proteins.

### 3.2 Protein structure alignment tools

Table 1 shows nine protein structure alignment tools for pairwise alignment, one structure library search tool that does not produce pairwise alignment, and four tools for multiple structure alignment. To the best of our knowledge, these are the most widely used protein structure alignment and search tools, with each being representative of its unique mechanism. We focus on pairwise structure alignment tools and library search tools that provide pairwise alignment patterns. We only perform multiple sequence/structure alignment on the phylogeny reconstruction task for a comprehensive comparison. More details on sources and implementations can be found in the Supplementary **Implementation and commands** section.

#### 3.2.1 DALI

DALI (13) utilizes the three-dimensional coordinates of each protein to calculate residue-residue (C*α*-C*α*) distance matrices. These matrices are decomposed into elementary contact patterns, such as hexapeptide-hexapeptide submatrices. Similar contact patterns in two matrices are paired and merged in to a set of these pairs. A Monte Carlo procedure is employed to optimize a similarity score defined in terms of equivalent intramolecular distances. By focusing on the agreement of intramolecular distances using an elastic similarity score variant, DALI accounts for the conformational flexibility of protein structures, which is a significant advantage over methods based on rigid-body superposition. DALI’s ability to handle sequence gaps, chain reversals, and free topological connectivity (non-sequential alignment) of aligned segments further enhances its versatility and accuracy in the classification of protein folds.

#### 3.2.2 TM-align

TM-align (14) (denoted as TMalign) is a widely used protein structure alignment method that employs the TM-score rotation matrix to determine the optimal superposition. This matrix is more sensitive to global topology than to local structure variations because the TM-score weighs residue pairs at smaller distances more heavily than those at larger distances. This weighting scheme is effective in capturing overall structural similarity. The consistency between using the TM-score rotation matrix for superposition and the TM-score similarity scoring function in dynamic programming allows for faster convergence and more accurate alignments than methods that use the RMSD-based rotation matrix.

#### 3.2.3 KPAX/KPAX-flex

A unique mechanism of KPAX is the use of Gaussian overlap scoring functions to compare backbone peptide fragments (17). By placing each C*α* atom in a canonical orientation, KPAX can quickly calculate the similarity of local backbone fragments without the need for computationally expensive least-squares fitting. This approach, combined with secondary structure-specific gap penalties, allows KPAX to produce tight overlays of secondary structure elements (SSEs) and accurate structural alignments.

#### 3.2.4 DeepAlign

DeepAlign (15) employs an iterative process similar to TMalign to determine the superposition but uses a complex scoring function that incorporates the following three components: TM-score, BLOSUM matrix, and internal coordinates (local similarity). The inclusion of the BLOSUM matrix and internal coordinates enables DeepAlign to capture local similarity and provide biologically meaningful alignments, albeit at the cost of increased running time.

For flexible alignment, KPAX-flex (22) assigns consecutive aligned residue pairs with distances less than 3Å as the first segment, forming a new alignment sub-problem. The remaining residues form new segments for separate alignments. Finally, all separate alignment patterns are merged to form the full flexible alignment pattern.

#### 3.2.5 US-align2

US-align2 (16) (denoted as USalign) can be used for non-sequential structural alignment with the fNS alignment setting. Unlike

TMalign, which uses residue-level TM-scores to calculate the score of a residue pair, US-align derives a score for a residue pair by calculating the distance between the 5th closest residues after superimposing the corresponding 5-residue-long fragments for this residue pair of the two proteins. An Enhanced Greedy Search (EGS) process then performs non-sequential alignment using the derived score matrix.

#### 3.2.6 DeepBLAST

DeepBLAST (7) utilizes ProtT5 to generate embeddings for each residue in a protein sequence. Unlike pLM-BLAST, DeepBLAST does not calculate cosine similarity directly between embeddings of two sequences. Instead, two convolutional neural networks are trained with TM-align alignment patterns as the ground truth in a differentiable dynamic programming framework (34) to generate match scores and gap scores, respectively. With this additional supervised learning process, DeepBLAST learns to capture structural similarity from sequence embeddings.

#### 3.2.7 TM-Vec

Published in the DeepBLAST paper (7), TM-Vec is a DL-based library search tool for large-scale structure database and it does not produce pairwise alignment patterns. It uses ProtT5 to generate embeddings the same as DeepBLAST, but a twin neural network is trained to generate two global embeddings for the two sequences in the alignment, so that the cosine similarity between the two embeddings predicts the TM-score value from TMalign.

#### 3.2.8 pLM-BLAST

pLM-BLAST (18) directly utilizes the pre-trained ProtT5 protein language model (pLM) to generate embeddings of all residues in a protein sequence. To calculate the score matrix for dynamic programming, it normalizes embedding vectors of residues across all dimensions and computes the cosine similarity between the normalized embedding vectors of a residue pair. These embeddings capture contextual information about each residue in a protein sequence, allowing for a more sensitive comparison of sequences than traditional methods. pLM-BLAST employs a prefilter step to enable fast library search for large databases, which treats sequence embeddings as 2D images and convolves one embedding with another to get cosine similarities of embedding slices of the two sequences.

#### 3.2.9 FoldSeek

FoldSeek (12) is a deep learning method that converts the original amino acid sequence into a new sequence with 20 letters (the 3Di alphabet), representing 20 states of interactions between a residue and its surrounding residues in the local conformation. Traits of the local conformation used to encode the original sequence includes distances and angles between a residue and its surrounding residues, forming a 10-dimension vector for each residue. Then a VQ-VAE encoder-decoder neural network model (35) is trained to convert each residue to one of the 20 states in the 3Di alphabet. The score matrix for the 3Di alphabet is estimated from a small set of the training data and then used for dynamic programming to determine the alignment pattern.

### 3.3 Structural overlap quality evaluation

To evaluate the alignment quality of structure alignment tools, we use the following three reference-independent metrics (14):

N align, RMSD, and TM-score. N align is defined as the number of aligned residue pairs between the two structures. RMSD stands for Root Mean Square Deviation, which measures the mean Euclidean distance between aligned residues of the two structures in the 3D coordinate system:

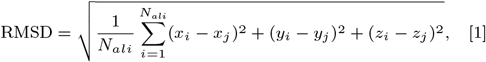

where *N*_*ali*_ is the number of aligned residues, and (*x*_*i*_, *y*_*i*_, *z*_*i*_) and (*x*_*j*_, *y*_*j*_, *z*_*j*_) are the coordinates of the paired residues from protein *i* and *j*, respectively.

Because the RMSD assigns equal importance to distances between all residue pairs, minor local structural deviations can lead to a high RMSD. In addition, the average RMSD for randomly associated proteins is influenced by the length of the structures compared, and higher N align often result in higher RMSD, which diminishes the interpretive value of the absolute RMSD value. However, a small N align could be insufficient for a biologically meaningful alignment pattern, even with a low RMSD. Therefore, the TM-score (36) was proposed to address the trade-off between N align and RMSD, which prefers a high N align and a low RMSD.

TM-score can be viewed as the length-normalized distance between two proteins:

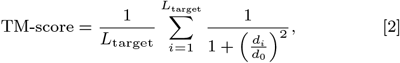

where *L*_target_ is the length of the target structure, *d*_*i*_ is the distance between the *i*-th pair of aligned residues, and *d*_0_ is a distance scale. The distance scale *d*_0_ is defined as:

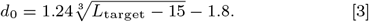

The target protein is defined as the shorter protein for alignment accuracy evaluation and phylogeny reconstruction. In homology detection and function inference, the target protein is defined as the query protein whose properties are to be determined by database search.

We also include the reference-dependent accuracy metric to assess the quality of pairwise alignment given the alignment patterns manually curated by human expects as the ground truth. These curated patterns are included in the Malidup and Malisam datasets. The accuracy of alignment is defined as the proportion of correctly aligned positions with respect to the given pattern.

### 3.4 Evaluation of homology detection accuracy

For the homology detection task, we follow the settings of the DALI benchmark pipeline (13), which queries 140 proteins over a dataset of 15211 proteins from the SCOP database (37). The target protein with the highest structural similarity score to the query protein is chosen as the reference, and the structural category label of the target protein defined by SCOP is transferred to the query protein. There are three hierarchical levels in the evaluation: family, superfamily, and fold, with increasing evolutionary distance and difficulty for alignment. At each level, homology detection is a binary classification task, where only protein pairs with the same category labels are treated as positive pairs. The pairwise alignment scores produced by alignment tools are used as probabilities to be positive pairs. The overall classification accuracy is defined as the maximum F1 score given a certain threshold at each level:

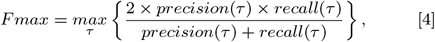

where *τ* is a flexible threshold to be determined for both recall and precision to obtain the maximum F1 score. Precision and recall for binary classification at each level are defined as:

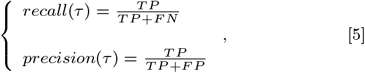

where TP is the number of true positive pairs, FN is the number of false negative pairs, and FP is the number of false positive pairs. A positive pair is determined when its alignment score is above the given threshold *τ*.

### 3.5 Evolutionary Tree Construction

Phylogeny reconstruction is to construct/predict an evolutionary tree based on quantitative distances across species. These distances can be inferred by either sequence or structure information. For pairwise methods such as TMalign and BLAST, metrics reflecting alignment qualities, including RMSD and TM-score, are recorded for distance matrix reconstruction. Specifically, we consider several identity metrics that utilize sequence and structure information: (1) Amino Acid Identity in structurally aligned regions (AA-Identity), (2) Amino Acid Identity of structurally aligned regions against the full length (AA-Identity-full), and (3) Spatial Overlap Identity (SO-Identity) that measures the percentage of structurally aligned positions over the full length of sequence. Both first two identities use the number of identical residues in structurally aligned regions as the numerator, but they use different denominators: AA-Identity uses the length of the aligned region as the denominator, whereas AA-Identity-full uses the full length of the sequence as the denominator. These two metrics utilize both sequence and structure information, which is different from the sequence identity in BLAST. SO-Identity is a metric based solely on structure.

All identity metrics and TM-score range from 0 to 1, thus we use 1*−metric* as the distance between any two proteins. For RMSD distance, we divide each RMSD value by the maximum RMSD of all pairs in the corresponding protein family so that all RMSD values are scaled between 0 and 1, and the scaled RMSD is used as the distance value. Following the workflow in Foldtree (6), we use the distance-based tool FastME (version 2.1.6.3) (38) to construct an evolutionary tree based on the distance matrix, and then set all negative branch lengths to 0. Then all trees are rooted using MAD (version 2.2) (39) in the default setting.

For multiple sequence/structure alignment (MSA) methods, we first retrieve the MSA pattern in the FASTA format from each tool and then use the FastTree (version 2.1.11) (40) to build the evolutionary tree. All trees are also rooted using MAD in the default setting.

### 3.6 Evolutionary Tree quality assessment

We use two metrics to assess the quality of the reconstructed evolutionary tree: the Robinson-Fould (RF) distance and the tree congruence score (TCS). The RF distance has been widely used as a measure to reflect the deviation of a predicted tree from a ground-truth tree, defined in terms of bipartition dissimilarity (41). The ground-truth tree for each protein family is included in the SwissTree database and this tree is constructed based on species phylogeny. The normalized-RF distance, ranging from 0 to 1, is calculated by dividing the original RF distance with the maximum number of all possible bipartitions of the trees being compared.

Following the Foldtree workflow, we also use the TCS score (42) as a reference-independent metric which does not require manually curated trees as the ground truth, but requires taxonomy labels of proteins retrieved from the UniProt database (43). The Foldtree version of the TCS score calculates the overlap of taxonomy labels between the two children of each internal node. The final TCS score is defined as the sum of TCS scores of all leaves.

The overlapping label set *s*(*x*) for each node is calculated as:

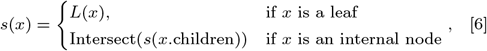

where *L*(*x*) is the length of taxonomy labels of each protein (leaf). Then the TCS score of a node is calculated as:

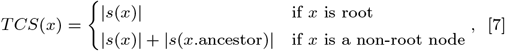

where |*s*(*x*)| is the length of the overlapping taxonomy label set of the corresponding internal node. The final TCS is defined as:

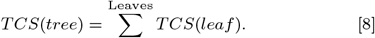

A normalized TCS score is reported by dividing the original TCS by the number of proteins in the corresponding family.

In summary, a low normalized RF distance or a high normalized TCS score indicate good quality of the predicted evolutionary tree.

### 3.7 Function inference accuracy evaluation

Protein function inference is a multi-class multi-label classification task. There are 677 GO functional labels for our filtered CAFA3-MF dataset, and each protein can be associated with more than one label. Fmax is the official metric of the CAFA competition and is defined the same as in equation [4]:

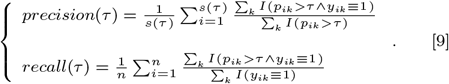

Here *s*(*τ*) denotes the number of proteins that are predicted with at least one function, *k* is the total number of labels and here *k* = 677, *p*_*ik*_ is the predicted score for the function, and *y*_*ik*_ is the ground truth with 1 indicating the existence of the function and 0 otherwise. *n* is the total number of proteins in the evaluation and here *n* = 1137.

### 3.8 Experiment setup

All experiments are conducted on a computer with an AMD EPYC 7302 CPU with 128G memory. All DL-based methods are run on a single RTX 3090 GPU with 24G memory.

## 4 Results

### 4.1 Alignment accuracy evaluation by reference-dependent and independent metrics

To objectively evaluate the alignment qualities of all methods, we focus on L align (the number of structurally aligned residual pairs), RMSD (averaged spatial distance in the aligned region), and TM-score (a length-normalized version of spatial distance). Fig. 2A illustrates that the flexible method KPAX-flexible (denoted as KPAX-flex) achieves the highest TM-score (0.69), indicating its superior structural alignment performance. This high TM-score for KPAX-flex is a result of its flexible alignment mechanism, producing a high L align of 63 and a notably lower RMSD of 1.92. TMalign, DeepAlign, and KPAX produce high TM-scores ranging from 0.51 to 0.52, with L align values between 58 and 61 and RMSD values ranging from 2.65 to 3.06. These traditional methods demonstrate good trade-offs between L align and RMSD, reflecting a preference for higher L align and lower RMSD. Between the two non-sequential methods DALI (non-open source version) and USalign, DALI produces alignments with higher RMSD, and USalign outperforms its sequential version TMalign on all three metrics. The three DL methods, pLM-BLAST, DeepBLAST, and Foldseek, tend to yield suboptimal superpositions with distinctly higher RMSD and lower TM-score than non-DL methods. Among the three, pLM-BLAST performs the worst, which could be due to its scoring matrix for dynamic programming being calculated from the cosine similarity between embeddings of a residue pair generated by the protein language model (pLM), and pLM may be unable to capture enough structure information. In comparison, DeepBLAST, a method trained to attain the alignment patterns from TMalign, and Foldseek, which encodes 3D structures into 1D strings, achieve the best DL-based performance. The baseline sequence-based alignment method, the Needleman-Wunsch (NW) algorithm, yields the worst RMSD and TM-score results among all methods although it provides longer alignment patterns. On reference-dependent evaluation, traditional sequence alignment method NW only attains an accuracy of 0.04 because Malisam is more difficult than Malidup (Fig. 2B). Methods differ more from each other in Malisam than in Malidup, with DALI, DeepAlign, and KPAX consistently performing well and DL-based methods performing poorly in Malisam. The alignment quality of Malidup has the same trend as that of Malisam across different methods but the difference in the overall performance is less distinguishable (Supplementary Fig. S1), because Malidup consists of duplicate units in proteins, whereas Malisam, containing known distant homologous pairs in different families, is more difficult for homology detection. We should be cautious that the ground-truth patterns we use from manual inspection are not necessarily correct but are used for reference only. Nevertheless, the alignment accuracy results still reflect how these tools differ in terms of the alignment patterns they determine.

**Fig. 2:**
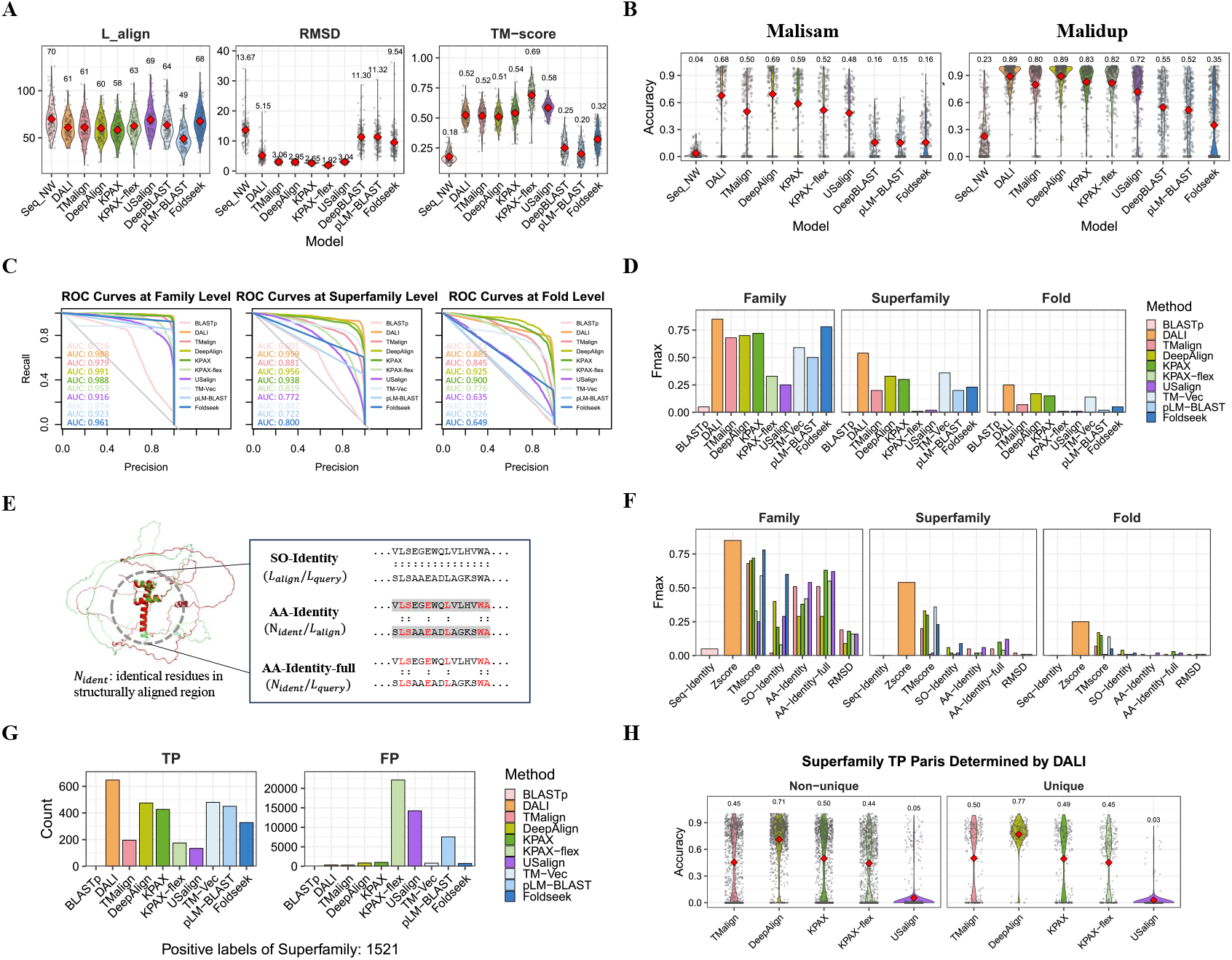
Performance of various structure alignment tools on structural overlap quality and homology detection. **A:** Reference-independent metrics on the Malisam dataset. Seq NW refers to the sequence alignment Needleman-Wunsch algorithm. **B:** Alignment accuracy evaluated on Malisam and Malidup datasets using provided manual alignment patterns. **C:** ROC curves for homology detection as a binary classification task at three different levels. **D:** Fmax scores of different structure alignment tools on the SCOP140 dataset. **E:** Illustration of three scoring metrics for measuring structure and sequence similarity. **F:** Fmax scores resulted from utilizing different scoring metrics. **G:** Counts of true positive and false positive pairs determined by different tools at the superfamily level of SCOP140. **H** : Alignment accuracies of different tools with respect to the DALI patterns of DALI-unique and -non-unique true possible pairs at the superfamily level.

### 4.2 Performance in homology detection

In Fig. 2C,D we find that DALI performs the best in overall homology detection accuracy, whereas KPAX-flex and USalign do not achieve comparable results despite their flexible and non-sequential mechanisms, respectively. DeepAlign and KPAX have comparable results across the three levels (family, superfamily, and fold). DL methods can deal with detection at the family and superfamily levels, but not at the fold level, which is harder than the other two levels. Foldseek, a fast DL library search tool for querying large structure databases, performs better than most traditional methods except DALI at the family level. TM-Vec, a DL tool that utilizes pLM to predict TM-score from a pair of sequences, has comparable performance with other traditional methods and outperforms most of them except DALI on the Fmax score at the superfamily level. pLM-BLAST can also handle family and superfamily level detection, with its fast comparison mechanism. Previous studies have found discrepancies between structural overlap performance and homology detection performance (7). Such an observation can also be seen here, as KPAX-flex and USalign produce the lowest Fmax scores at all three levels, whereas these two methods yield the best superpositions with the highest TM-scores in the Malisam dataset. The homology detection accuracy is more consistent with alignment accuracy than that with TM-score. However, although KPAX-flex and USalign have comparable performance in accuracy with TMalign based on manually aligned patterns, these two methods do not exhibit the same level of performance of TMalign in terms of Fmax score at all three levels. As expected, sequence-based alignment BLASTp cannot handle the structure-based homology detection task well, showing a low accuracy. Compared to previous studies, we are the first to perform a systematic comparison in the SCOP database across different methods.

To further explore the utilization of structure and information, we consider three ways of utilizing structure information: SO-Identity, AA-Identity, and AA-Identity-full (Fig. 2E). SO-Identity, referred to the SO (spatial overlap) metric in other studies (44), is the percentage of structurally aligned length over the full length of the query protein. AA-Identity is defined as the percentage of identical residues in the structurally aligned region, and AA-Identity-full is similar to AA-Identity but over the full length of the query protein instead of the length of the structurally aligned region. Other structure-based metrics such as TMalign, RMSD, and Zscore (utilized only by DALI) are also included for comparison. Methods including DeepAlign, pLM-BLAST, and Foldseek also incorporate sequence information in their scoring functions. In Fig. 2F we find that Zscore, generated from DALI 2D distance matrix representation, performs the best. Among other traditional methods that are based on the 3D coordinate representation, the structure-based metric TM-score is more suitable for homology detection, whereas SO-Identity, AA-Identity, and AA-Identity-full do not produce comparable results. Fig. 2G shows that the flexible method KPAX-flex and the non-sequential method USalign, and a DL method pLM-BLAST introduce many false positive pairs. The reason for these performance differences lies in the mechanisms of these methods, especially in how they match small proteins to large multiple-domain proteins. In addition, the scoring function plays an important role. The value of structure similarity can be more important than the alignment pattern, and the detection accuracy can be less related to the alignment accuracy than previously believed. We assess the difference between alignment patterns produced by DALI and those by others. We use the former as the ground truth since DALI finds out the highest number of true positive pairs, and then calculate the alignment accuracy of other tools with respect to the ground-truth pattern. We only show results at the superfamily level, because previous studies define protein pairs sharing the same superfamily but falling in different families as distant homology (37). We find that although DALI determines more than 100 true positive pairs (Fig. 2H), the alignment accuracies of DALI-unique true positive pairs attained by other tools are not necessarily lower than those of non-DALI-unique pairs. Results at family and fold levels are shown in Supplementary Fig. S2. These results suggest that even though different tools may provide similar alignment patterns, the scoring functions they use to produce the similarity values could make a difference in the performance of homology detection and other tasks.

### 4.3 Evaluation of phylogeny reconstruction

To evaluate how well different tools perform in reconstructing evolutionary relationships across species, we first perform pairwise alignment using all traditional non-DL methods and two DL methods pLM-BLAST and DeepBLAST. With BLASTp and Foldseek, we perform exhaustive searches to retrieve all vs. all comparison results. We also include several multiple alignment tools such as sequence-based Clustal Omega (denoted as Seq ClustalO) and structure-based mTMaligna and 3DCOMB for a comprehensive comparison. Results show that, in general, multiple alignment tools predict the most accurate evolutionary trees with consistently lower normalized RF distances and higher normalized TCS scores than other pairwise methods except Foldseek (Fig. 3A). In addition, Foldmason, the multiple alignment version of Foldseek, yields the second lowest normalized RF distance, slightly worse than Seq ClustalO. DALI performs the worst in both RF distance and TCS score, and for other trees are shown in Supplementary Fig. S7). The number of consistent columns reflects the difficulty of phylogeny reconstruction for a tree (Supplementary Table S3).

**Fig. 3:**
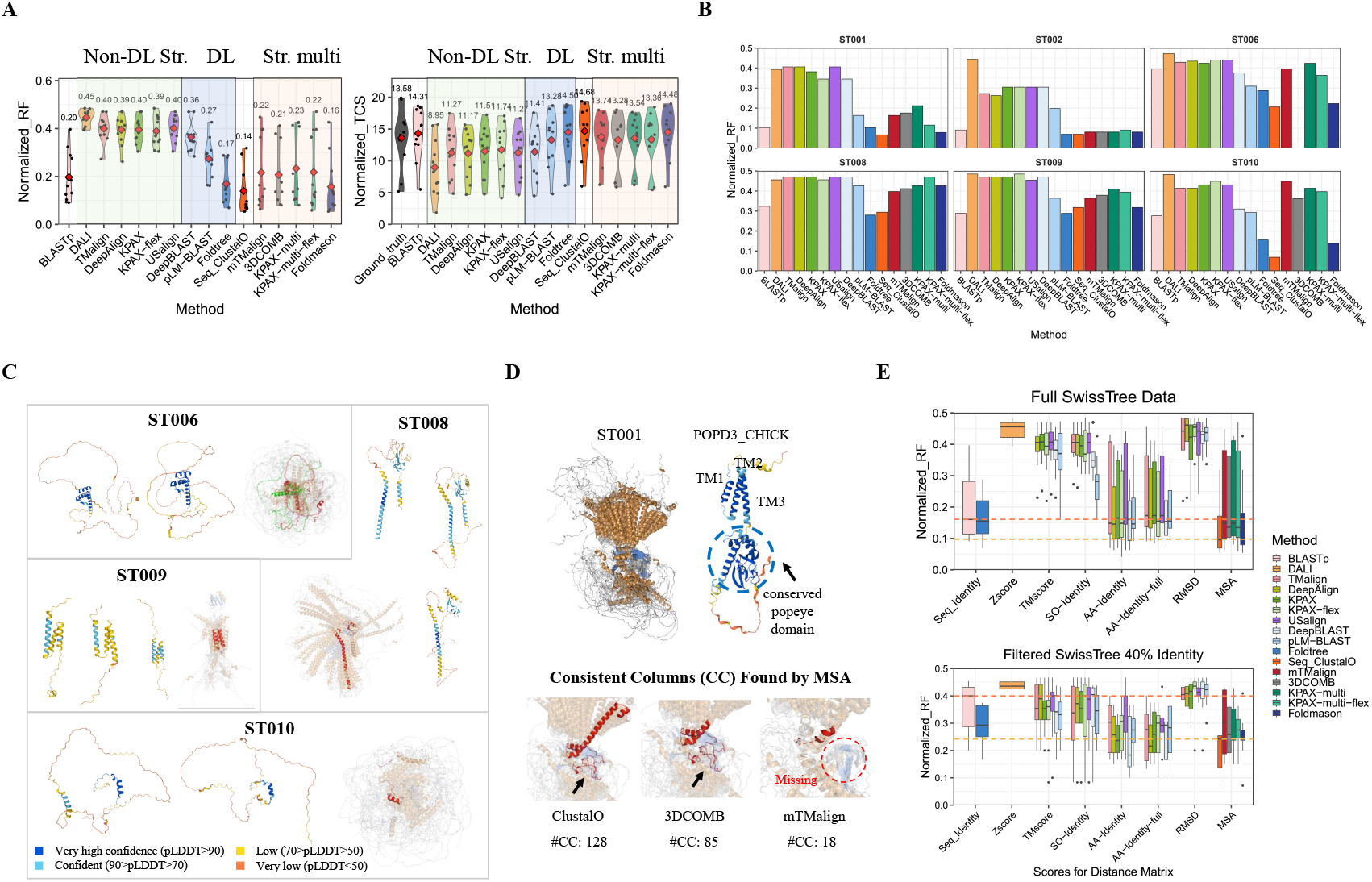
Performance of nine structure alignment tools on phylogeny reconstruction. **A:** RF distances and TCS scores of different tools on trees ST001-ST010 of SwissTree database. **B:** RF distances of different tools on each tree. **C:** Four protein families whose structures are hard to predict. **D:** Multiple structure alignment patterns of ST001. **E:** RF distances resulted from different tools under different sequence identity filtering conditions. The pink and orange dashed lines indicate the normalized RF levels of BLASTp and Seq ClustalO, respectively.

Our results seem to suggest that traditional structure-based methods are unsuitable for reconstructing phylogeny across species. To further investigate whether incorporating sequence information into structural information can help, we assess the effects of a wide range of scores for the distance matrix on RF distances. In the top panel of Fig. 3E, we find that normalized RF distances from AA-Identity of traditional structure alignment tools and the two DL methods are lower than those from BLASTp, and are higher than those from multiple alignment methods. The performance of using the two sequence-structure-combining metrics AA-Identity and AA-Identity-full is better than that of using sequence-based metrics Zscore, TM-score, SO-Identity, and RMSD. In particular, both RMSD and Zscore produce significantly worse results than the others. This result suggests that structure information is useful to further improve the accuracy of predicting evolutionary trees over using only sequence information. When we filter the SwissTree database to retain sequences with identities lower than 40%, structural information becomes more useful than sequence information, as TM-score and RMSD outperform sequence identity by BLASTp on RF distances.

This phenomenon does not appear on the full SwissTree dataset. In addition, DeepAlign, KPAX and DeepBLAST achieve the best performance when using both structure and sequence information, better than Seq ClustalO (attained by DeepBLAST and KPAX using AA-Identity, and DeepAlign using AA-Identity-full). When sequence identities across proteins in a family are lower than 40%, there are fewer orthologs but more paralogs with highly different structures and functions in this family. Such big differences are not pivotal for detecting close homology on the full dataset. These results suggest that as sequence identities across proteins in a protein family become lower, especially below the twilight zone (*<*25%), structure alignment is better than sequence alignment to recover the evolutionary relationship in the protein family.

### 4.4 Accuracy evaluation of function inference

Protein function inference is a difficult task that requires information from different sources such as sequences, structures, interaction networks, and documents. Utilizing structure data for homology search has not been comprehensively explored due to the lack of structure data and fast search tools. With these two challenges tackled by recent technology advances, we compare the performance of structure alignment methods as homology search strategy with traditional sequence alignment methods on the protein function inference task. Several alignment scores such as RMSD, TM-score, and SO-Identity are evaluated (Supplementary Fig. S8). We only report the highest Fmax score generated by using TM-score or using SO-Identity for each tool. With TMalign and USalign, the TM-score gives higher Fmax scores. With DeepAlign, KPAX, and KPAX-flex, the SO-Identity produces better performance. For BLASTp, pLM-BLAST, and Foldseek, since they are library-searching tools with rankings, we use their default ranking scores, which are local sequence identity, embedding cosine similarity, and Fident (sequence identity defined in the 3Di alphabet used in Foldseek), respectively, for evaluation. Results in Fig. 4A show that all methods included in our function reference benchmark have comparable performance except USalign. As DALI fails to preprocess many proteins in AlphaFoldDB, we do not include results from DALI. The top three alignment methods based on Fmax scores are KPAX, TM-Vec, and Foldseek for function inference in the CAFA3-MF setting. Further investigation on scoring metrics shows that the AA-Identity-full score generally produces higher Fmax scores for traditional pairwise alignment tools than that from BLASTp (Supplementary Figure. S8).

**Fig. 4:**
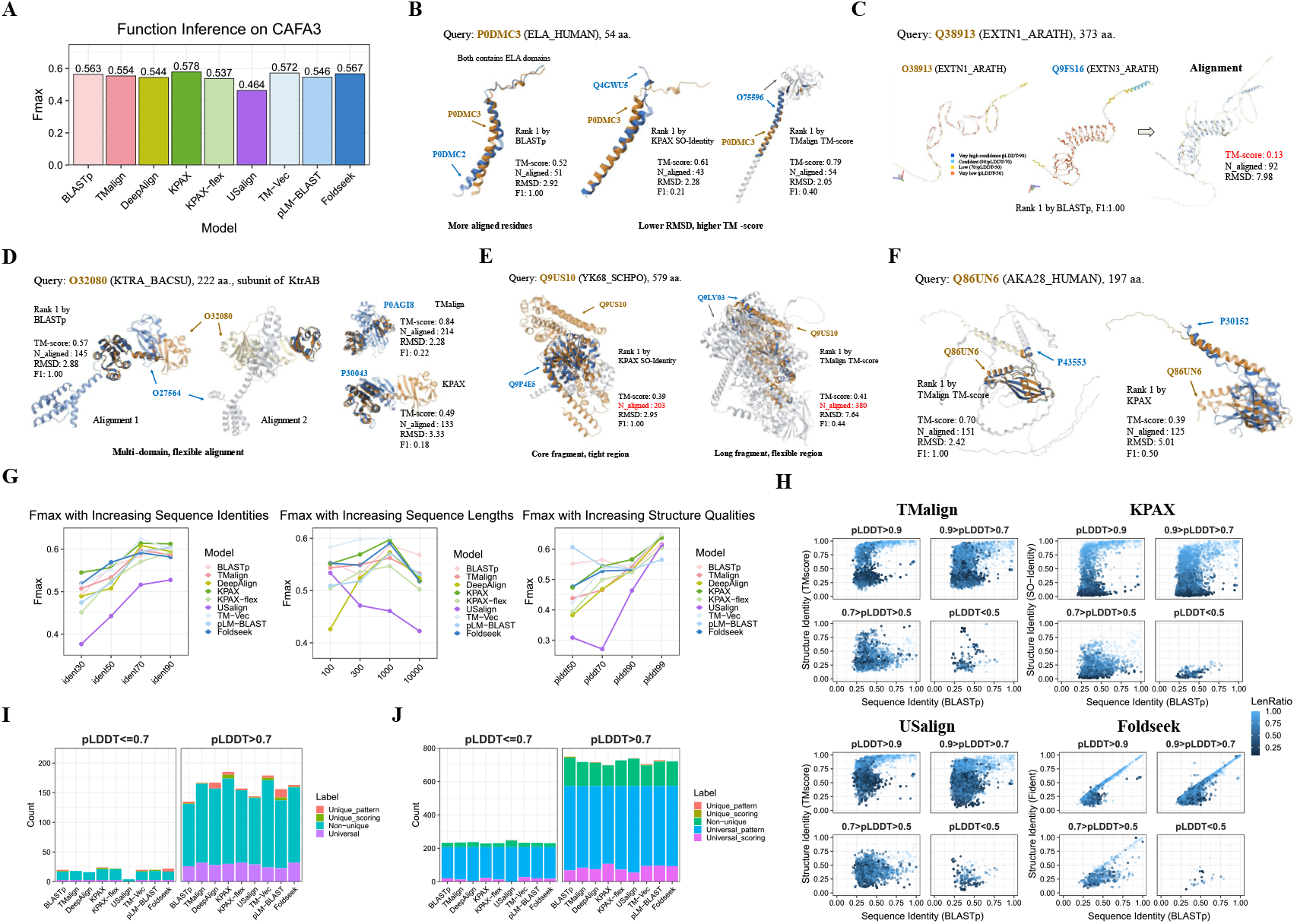
Performance of nine structure alignment tools on function inference. **A:** Fmax scores on the CAFA3-MF dataset. **B:** Structural alignment visualization of P0DMC3 P0DMC2, P0DMC3 Q4GWU5, and P0DMC3 O75596. **C:** Structural alignment visualization of O38913 Q9FS16. **D:** Structural alignment visualization of Q9US10 Q9P4E5 and Q9US10 Q9LV03. **E:** Structural alignment visualization of Q86UN6 P43553 and Q86UN6 P30152. **F:** Structural alignment visualization of O32080 O27564, O32080 P0AGI8, and O32080 P30043. **G:** Trends of Fmax scores under different conditions of sequence identities, query lengths, and AlphaFold predicted structure qualities. **H:** Scatter plots of sequence similarity versus structure similarity of different tools. **I:** Distribution of hits (targets with F1=1.0) recovered by different tools. **J:** Distribution of targets missed by different tools.

It is surprising that BLASTp, widely used for homology search strategy in function inference, produces an Fmax score similar to the solely structure-based method KPAX and the structure-sequence combined method Foldseek. Although it is commonly believed that structure is more conserved than sequence (3) and thus structure similarity is supposed to better infer function conservation than sequence similarity, this belief has not been extensively tested in a large database in a real-life setting. In contrast, our results indicate that whether utilizing sequence information or structure information makes little difference as the homology search strategy in the performance of function inference. To investigate the reason and further illustrate how different tools determine the best aligned structures, we manually inspect superpositions of several target proteins determined by different tools on the same query protein. In Fig. 4B, the function similarity between query protein P0DMC3 and target protein P0DMC2 is recovered by BLASTp, indicated by an F1 score of 1.0 between GO terms of the two proteins. While the two target proteins found by structure alignment methods KPAX and TMalign have higher TM-scores, their corresponding F1 scores are low. This phenomenon occurs because when the structure of the query protein is simple (short and with few secondary structures), structure alignment is less likely to reveal functional similarity due to the loss of information at the amino acid level. Current structure alignment methods rely on the mapping between two proteins on the positions of the central carbon atoms for each corresponding pair of residues, but they discard side chain information, resulting in missing physicochemical information that is reflected by types of amino acids. In addition, BLASTp is insensitive to the quality of AlphaFold predicted structures, as illustrated in Fig. 4C, and has relatively high performance on proteins with pLDDT scores less than 50, as shown in Fig. 4G. Furthermore, sequence alignment can deal with flexible alignment, as shown in 4D, whereas rigid methods including TMalign and KPAX can only map part of the query protein to the target protein. These observations indicate that sequence alignment is crucial and unreplaceable by structure alignment for function inference.

The mechanism to handle partial or local alignment is one of the main factors that contribute to the performance of structure alignment tools. It is hard to determine the best superposition that aligns the biologically related regions when matching a small protein to part of a large protein, or vice versa. Finding the most biologically related regions of two proteins must consider both L align and RMSD. TMalign, as a global method, tends to yield longer alignments with higher RMSD, whereas KPAX, which takes local conformation into account, generates more conservative and compact alignment patterns with smaller RMSD (Fig. 4E). KPAX’s local indexing mechanism to deal with local conformation makes KPAX outperform other global structure alignment methods in function inference. However, the trade-off between L align and RMSD becomes more complicated as the query and target lengths differ more, and the conservative alignments produced by KPAX do not necessarily give better results than TMalign (Fig. 4F).

To further analyze how the performance of these tools is affected by different factors, we categorize query proteins based on their sequence identities shared with target proteins, sequence lengths, and pLDDT scores. In the left panel of Fig. 4G, all methods show an increasing trend on Fmax scores as sequence identities increase from 30% to 70%, and canonical structure alignment methods including TMalign, DeepAlign, and KPAX, as well as TM-Vec and Foldseek, encounter a turning point at identity values around 90. One possible explanation is that those proteins are highly similar to their family members in the dataset and they are almost identical in structures. The evolutionary traits that distinguish them are better captured by methods that greatly utilize sequence information such as BLASTp and pLM-BLAST. In the middle panel, all methods except USalign show an increasing trend on Fmax scores as sequence lengths increase up to 1000 aa., and encounter significant drops of performance after 1000 aa., probably due to the partial alignment issue. Result of sequences greater than 1000 aa. is unavailable for TM-Vec because we exclude sequences that exceed 1000 aa. to ensure that the TM-Vec computation does not surpass our GPU memory resource. In the right panel, there are trends of increasing Fmax scores with increasing pLDDT in general. The two methods that utilize sequence information, BLASTp and pLM-BLAST, exhibit performance drop when their pLDDT scores increase from 50 to 70, and they perform the best when the pLDDT level is low. Foldseek, which takes pLDDT into account for ranking, can handle the quality problem relatively well.

To further illustrate the effects of sequence identity, sequence length, and pLDDT score, especially partial alignment, we analyze the correlation between structure similarity scores and sequence similarity scores (identity) in these tools. We only plot the top 10 target proteins from BLASTp for each query protein. As shown in Fig. 4H, protein pairs with lower length ratios (greater differences in lengths) are painted with darker blue, thus high sequence identities in the x-axis imply local structure alignment patterns for these pairs. First, plateaus appear as sequence identity increases, suggesting that when sequence identity is low, the structure similarity can be high enough to reveal functional relation. A clear linear correlation is seen only when structure quality is high (pLDDT*>*0.9). Second, KPAX tends to produce lower structure similarity than TMalign and other tools for partial alignment, reflected by lower y-axis positions of darker points, showing its preference for alignment with similar lengths and thus conserved patterns. In contrast, global methods such as TMalign, DeepAlign, and USalign tend to produce alignment patterns with much higher structure similarities, which may return target proteins with little biological relation. The difference between KPAX and global methods has little to do with the choice of scoring methods (Supplementary Fig. S9). Interestingly, Foldseek exhibits a nearly perfect linear correlation between sequence identity and its Fident score. This suggests that Foldseek performs more similarly to sequence alignment than structural alignment. Foldseek, which encodes 3D structures into 1D strings based on local conformation, is thought to be a combination of sequence and structure alignment.

As shown in Fig. 4I, we define a target protein with a 1.0 F1 score with respect to the query protein as a hit. Hits recovered by only one method are denoted as **Unique**, and those recovered by all methods are denoted as **Universal**. The rest are marked as **Non-unique**. We further consider whether these methods produce a similarity score high enough for a functionally related target protein. If a unique protein is within the top-10 lists of at least two other methods, we denote it as **Unique scoring**, meaning that such a hit is recovered attributed to the scoring method that rates it higher than other similar target proteins. Otherwise, we suppose the hit is recovered owing to a method’s ability to determine the biologically meaningful pattern, e.g., when encountering a partial alignment situation (**Unique pattern**). We find that high-quality query proteins are more likely to be unanimously inferred, and DeepAlign, KPAX, TM-Vec, and pLM-BLAST could be useful in dealing with some special cases regarding ranking homologous candidates and partial alignments, given their higher counts of unique hits than others. In Fig. 4J we use similar definitions for **Universal** and **Unique** in missing targets, but we define **Universal scoring** and **Unique scoring** as those of which the missing targets are within the top-10 lists. We find that most missing targets are unanimously missed, and that their missing is largely due to difficult matching cases such as partial alignment. These observations exist for both low- and high-quality proteins.

### 4.5 Performance comparison on large-scale database queries

We use CAFA3-MF, a dataset in the Kaggle competition, to study the performance of querying large databases. CAFA3-MF contains 1137 query proteins and 32421 target proteins. These queries make 30 million pairwise comparisons, which will take months for TMalign to finish querying on a single-core CPU. We summarize our test results in Table 2. The execution time is measured with the built-in /usr/bin/time function for only the querying or searching command of a certain tool but does not include time spent on library building. All pairwise structure alignment tools take a long time to traverse the entire target database, with KPAX running the fastest, and DeepAlign, a method with a complex scoring function, taking the longest time. Library search methods run much faster than pairwise alignment tools, attributed to their indexing and pre-filtering techniques, especially those utilizing the GPU, such as TM-Vec and Foldseek. We group query proteins based on their lengths to see how each tool scales to query length. Results (Fig. 5A) show that pairwise alignment methods exhibit quadratic trends on both running time and memory consumption with the increase of protein length. DeepAlign is the slowest due to its scoring function incorporating both local and global information, and KPAX-flex consumes the most CPU memory to deal with various possible conformation. USalign, a non-sequential method, also requires high CPU memory and long execution time. Most of the library search methods in Fig. 5B, except GTalign, show linear trends with the increase of length of query proteins except GTalign. TM-Vec and pLM-BLAST have a drop in the execution time with query length increasing from less than 100 aa. to 100-300 aa. because additional time is taken to load models into GPU and process the library and there are more proteins with a length of 100-300 aa. than those with a length less than 100 aa. (Supplementary Table S4). GTalign, although utilizing the GPU and other accelerating techniques, operates in the 3D coordinate system and does not use DL architectures, resulting in the slowest performance but the least memory consumption as a library search tool. In contrast, DL methods, such as TM-Vec and pLM-BLAST, encode the structural information as model parameters of their DL architectures, consuming more than 10 GB of CPU/GPU memory. These results suggest that, although pairwise methods such as TMalign and KPAX generally perform well in downstream tasks, DL methods could be more suitable for tasks that involve a large number of queries (millions of comparisons) such as homology detection and function inference. Acceleration of traditional methods such as DALI, KPAX, and DeepAlign with advanced indexing, filtering, and paralleling techniques is also a promising direction to facilitate the application of traditional methods to tasks that require intensive pairwise comparisons, as exemplified by the success of GTalign compared to its CPU predecessor, TMalign.

**Table 2.** Comparison of different tools in terms of input type, hardware, strategy, runtime, and memory, estimated with one CPU or one 3090 GPU on 36 millions queries of CAFA3-MF. * denotes that we exclude sequences that exceed 1000 aa. to ensure that the TM-Vec computation does not surpass the available GPU memory resource.

| Model | Input |  | Hardware |  | Strategy |  | Runtime | Memory |
| --- | --- | --- | --- | --- | --- | --- | --- | --- |
|  | Sequence | Structure | CPU | GPU | Pairwise | Search |  |  |
| BLASTp | ✓ |  | ✓ |  |  | ✓ | 436 sec. | 60 MB |
| Diamond | ✓ |  | ✓ |  |  | ✓ | 2.4 sec. | 162 MB |
| TMalign |  | ✓ | ✓ |  | ✓ |  | ~470 days | 14.2 MB |
| GTalign |  | ✓ |  | ✓ |  | ✓ | 5258 sec. | 630 MB |
| DeepAlign |  | ✓ | ✓ |  | ✓ |  | ~2800 days | 52.3 MB |
| KPAX |  | ✓ | ✓ |  | ✓ |  | ~290 days | 956 MB |
| KPAX-flex |  | ✓ | ✓ |  | ✓ |  | ~335 days | 956 MB |
| USalign |  | ✓ | ✓ |  | ✓ |  | ~791 days | 277 MB |
| TM-Vec | ✓ |  |  | ✓ |  | ✓ | *148 sec. | 16691 MB |
| pLM-BLAST | ✓ |  |  | ✓ |  | ✓ | 720 sec. | 19456 MB |
| Foldseek |  | ✓ |  | ✓ |  | ✓ | 51 sec. | 3277 MB |

**Fig. 5:**
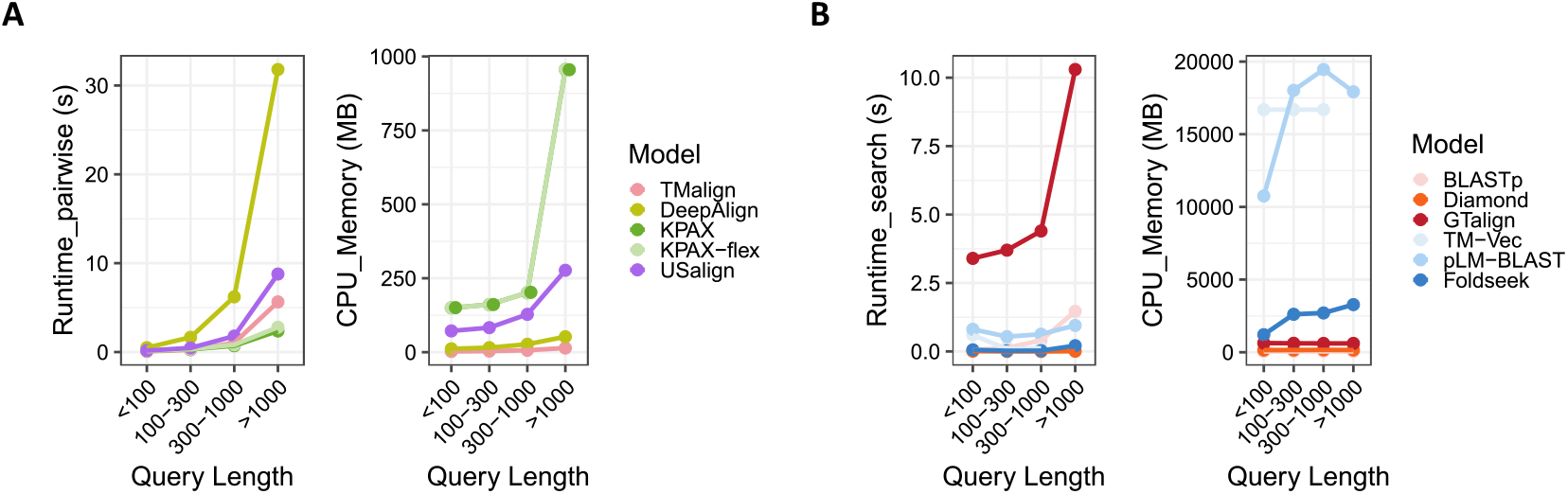
Performance of eleven tools on large-scale querying on the CAFA3-MF dataset. **A:** Running time and memory consumption of pairwise alignment methods. **B:** Running time and memory consumption of library search methods.

## 5 Discussion

In this study, we assess the performance of nine existing structure alignment tools in three downstream tasks. As previous studies found discrepancies between structural overlap quality and homology detection performance, we not only further confirm this observation in our homology detection benchmark, but also show that the performance of predicting evolutionary tree and protein function is less correlated to structural overlap quality than commonly believed. For example, KPAX-flex and USalign, the two methods that produce the largest structural overlaps and achieve the highest TM-scores, perform relatively worse than others in downstream tasks because the alignment patterns they determine are less biologically meaningful than those from the other tools, as addressed in a study (46). Nevertheless, flexible methods and non-sequential methods can still be useful to handle special structure alignment scenarios. In addition to alignment patterns, scores generated by structural overlap metrics, such as Zscore, TM-score, and RMSD, are important for several downstream tasks, as their accuracies are sensitive to the values of similarity measurement. The effect of scoring is demonstrated by a wide range of accuracy values using different scoring metrics in the three downstream tasks. There is no definite winner method for all tasks. On homology detection, TM-score produces the best results and RMSD performs the worst. On phylogeny reconstruction, TM-score and SO-Identity have comparable performance on both original and low-identity datasets. On function inference, SO-Identity is better for those considering local conformation such as DeepAlign and KPAX, and TM-score is better for global alignment methods such as TMalign and USalign. These observations suggest that an evolution-aware scoring metric may help produce a better ranking for evolution-related tasks.

We are interested in whether structure information is more useful than sequence information in downstream tasks, and how to better utilize both kinds of information. We introduce AA-Identity and AA-Identity-full, both of which only consider residue equivalence in structurally aligned regions. On homology detection, which focuses on structural superposition, sequence alignment yields extremely low accuracy, and the introduction of sequence information using AA-Identity does not improve the performance. In contrast, sequence alignment produces consistently good results on phylogeny reconstruction, which is previously handled by sequence-based methods only. Although structure information alone cannot accurately reconstruct the hierarchy of closely homologous proteins, we show that using AA-Identity can greatly improve the accuracy of predicted evolutionary trees by traditional structure alignment methods. Foldseek, whose mechanism are found similar to sequence alignment, has a comparable performance to BLASTp. We also find that when sequence identity is low, structure alignment becomes more useful than sequence alignment, and in some cases better than multiple alignment. We believe this advantage can become larger when the identity is lower than 30%, as observed in a study (26). In such a case, the phylogenetic tree is rather a function tree than a species tree. For function inference, structure alignment methods have comparable results to sequence alignment methods, and using AA-Identity-full further improves the Fmax scores of all traditional structure alignment methods. All these results suggest that sequence alignment is still crucial for downstream tasks, and combining sequence information with structure information could be useful in certain scenarios. What is the best way to utilize both sequence and structure information is an open problem.

Partial/local alignment is another major challenge in tasks such as homology detection and function inference where multi-domain proteins are present. When large query proteins are matched to small target proteins in the database, only part of the entire realm of functions is recovered depending on what functional region is aligned. When small query proteins are matched to large target proteins, we need to decide what region is the most biologically related. These problems require further investigation to resolve. One direction is to create databases for functional structure elements/domains that can be seen as smallest structural units of functions. Another solution is to utilize DL methods.

Previously it was difficult to evaluate phylogeny reconstruction and function inference with structure alignment due to the lack of structure data. In this study, we use predicted structures in AlphaFoldDB for all proteins in the two tasks of our benchmarking study. Because predicted structures can be incorrect, the quality of these data, reflected by the pLDDT score, is a crucial factor to assess. For phylogeny reconstruction, protein families with low pLLDT scores are harder to accurately reconstruct their evolutionary trees. Low pLLDT scores also result in decreasing accuracy of all methods in function reference. However, the low quality of some predicted structures is not necessarily considered as a flaw of the AlphaFold model itself. The pLDDT score also reflects the flexibility and irregularity of a protein. Proteins with loose/flexible regions are intrinsically hard to align due to a variety of possible conformations in the 3D space, as seen in the SwissTree ST006 family. In such cases, sequence alignment is not affected by the quality of predicted structures. Therefore, tasks with many proteins of low pLLDT scores could be better tackled by sequence-based and DL methods.

In general, TMalign is still the canonical choice for a good balance between execution time and memory consumption, but KPAX outperforms TMalign in many cases and runs 2x faster at the cost of larger memory consumption. All traditional pairwise structure alignment methods are unsuitable for large-scale queries due to the unacceptable long running time quadratic to query length, except for the recent GTalign, a GPU version of TMalign. Foldseek, as a combination of sequence alignment and structure alignment, is promising in both speed and accuracy in downstream tasks. Our results suggest that its similar mechanism to sequence alignment could be the reason that it performs significantly better than other non-multiple structure alignment tools on the phylogeny reconstruction task. Its ability to characterize 3D structure information into 1D strings is demonstrated by relatively high accuracy on the homology detection task. In addition, Foldseek, a library-searching tool, ranks the alignments by combining the default metric Fident and the pLDDT score. This mechanism helps Foldseek alleviate the influence of proteins with low AlphaFold prediction quality. In summary, we recommend both TMalign and KPAX for most tasks because of their good balance between running time and memory consumption, and relatively good and stable accuracy performance in downstream tasks. In tasks that require a large number of pairwise comparisons, such as homology detection and function inference, Foldseek is the best choice to attain a good result within a reasonable time. Although TM-Vec produces results comparable to Foldseek in the three tasks, it takes significantly longer to process sequence data than Foldseek, including model loading and embedding generation using pLM (data not shown). The trade-off between task accuracy and speed remains the major consideration in developing new alignment for downstream tasks.

We acknowledge the following limitations in our study. First, although we focus on evaluating protein structure alignment tools in various scenarios, some scenarios cannot be properly resolved by structure alignment methods. (1) Regions in a protein could differ in volutionary rates, and evolutionary traits that reflect important biological relationships could reside in variable loop regions, as shown in a study on Covid evolution (45). In such a case, using sequence alignment is difficult to determine what part of the protein is more important than the rest in a specific scenario. In addition, as pointed out in a previous study (47), structure alignment cannot capture discrete evolutionary events in amino acid sequences as well as the sequence alignment, due to the highly variable conformation. (2) The size and comprehensiveness of the database affect the accuracy of protein function inference. Databases with a wide range of representative structures and functions result in better performance. We also find that annotations are missing in both the query and target datasets that we use in MAFA3-MF, but this issue can be alleviated by incorporating the updated information in UniProt. (3) Alignment-based methods are unlikely the best choice for function inference. Alignment usually takes a long time, especially for traditional pairwise methods. In such a case, using DL methods would be of great benefit. Most of the models with top performance for the recent CAFA5 competition use DL architectures. Their authors find that the DL components are crucial for a high performance and outperform the alignment components. However, using alignment as a homology search strategy improves the performance of an ensemble model.

## 6 Data availability

Codes are available at https://github.com/georgedashen/StructAlign. Data is provided on https://zenodo.org/records/14938229.

## 7 Supplementary data

Supplementary Data are available at NAR Online.

## 8 Acknowledgements

Z. C. collects data, conducts experiments, and writes the manuscript. X. Z. helps install several tools and provides insight in mechanism explanation of tools, result analysis and discussion. W. Y. helps in discussing case study and result interpretation. Q. L. supervises this study, provides guidance on result analysis and interpretation, and revises the manuscript.

## References

1. Zhang, C., Shine, M., Pyle, A. M., and Zhang, Y. (2022) US-align: universal structure alignments of proteins, nucleic acids, and macromolecular complexes. Nat. Methods, 19(9), 1109–1115.

2. Berman, H. M., Battistuz, T., Bhat, T. N., Bluhm, W. F., Bourne, P. E., Burkhardt, K., Feng, Z., Gilliland, G. L., Iype, L., Jain, S., et al. (2002) The protein data bank. Acta Crystallogr., Sect. D: Biol. Crystallogr., 58(6), 899–907.

3. Illergård, K., Ardell, D. H., and Elofsson, A. (2009) Structure is three to ten times more conserved than sequence—a study of structural response in protein cores. Proteins: Struct., Funct., Bioinf., 77(3), 499–508.

4. Petrey, D., Fischer, M., and Honig, B. (2009) Structural relationships among proteins with different global topologies and their implications for function annotation strategies. Proc. Natl. Acad. Sci. U. S. A., 106(41), 17377–17382.

5. Carpentier, M. and Chomilier, J. (2019) Protein multiple alignments: sequence-based versus structure-based programs. Bioinformatics, 35(20), 3970–3980.

6. Moi, D., Bernard, C., Steinegger, M., Nevers, Y., Langleib, M., and Dessimoz, C. (2023) Structural phylogenetics unravels the evolutionary diversification of communication systems in gram-positive bacteria and their viruses. bioRxiv, pp. 2023–09.

7. Hamamsy, T., Morton, J. T., Blackwell, R., Berenberg, D., Carriero, N., Gligorijevic, V., Strauss, C. E., Leman, J. K., Cho, K., and Bonneau, R. (2024) Protein remote homology detection and structural alignment using deep learning. Nat. Biotechnol., 42(6), 975–985.

8. Ogden, T. H. and Rosenberg, M. S. (2006) Multiple sequence alignment accuracy and phylogenetic inference. Syst. Biol., 55(2), 314–328.

9. Lee, D., Redfern, O., and Orengo, C. (2007) Predicting protein function from sequence and structure. Nat. Rev. Mol. Cell Biol., 8(12), 995–1005.

10. Jumper, J., Evans, R., Pritzel, A., Green, T., Figurnov, M., Ronneberger, O., Tunyasuvunakool, K., Bates, R., Žídek, A., Potapenko, A., et al. (2021) Highly accurate protein structure prediction with AlphaFold. Nature, 596(7873), 583–589.

11. Varadi, M., Bertoni, D., Magana, P., Paramval, U., Pidruchna, I., Radhakrishnan, M., Tsenkov, M., Nair, S., Mirdita, M., Yeo, J., et al. (2024) AlphaFold Protein Structure Database in 2024: providing structure coverage for over 214 million protein sequences. Nucleic Acids Res., 52(D1), D368–D375.

12. Van Kempen, M., Kim, S. S., Tumescheit, C., Mirdita, M., Lee, J., Gilchrist, C. L., Söding, J., and Steinegger, M. (2024) Fast and accurate protein structure search with Foldseek. Nat. Biotechnol., 42(2), 243–246.

13. Holm, L., Laiho, A., Törönen, P., and Salgado, M. (2023) DALI shines a light on remote homologs: One hundred discoveries. Protein Sci., 32(1), e4519.

14. Zhang, Y. and Skolnick, J. (2005) TM-align: a protein structure alignment algorithm based on the TM-score. Nucleic Acids Res., 33(7), 2302–2309.

15. Wang, S., Ma, J., Peng, J., and Xu, J. (2013) Protein structure alignment beyond spatial proximity. Sci. Rep., 3(1), 1448.

16. Zhang, C. and Pyle, A. M. (2022) A unified approach to sequential and non-sequential structure alignment of proteins, RNAs, and DNAs. iScience, 25(10).

17. Ritchie, D. W., Ghoorah, A. W., Mavridis, L., and Venkatraman, V. (2012) Fast protein structure alignment using Gaussian overlap scoring of backbone peptide fragment similarity. Bioinformatics, 28(24), 3274–3281.

18. Kaminski, K., Ludwiczak, J., Pawlicki, K., Alva, V., and Dunin-Horkawicz, S. (2023) pLM-BLAST: distant homology detection based on direct comparison of sequence representations from protein language models. Bioinformatics, 39(10), btad579.

19. Sievers, F. and Higgins, D. G. (2014) Clustal Omega, accurate alignment of very large numbers of sequences. Multiple Sequence Alignment Methods, pp. 105–116.

20. Dong, R., Peng, Z., Zhang, Y., and Yang, J. (2018) mTM-align: an algorithm for fast and accurate multiple protein structure alignment. Bioinformatics, 34(10), 1719–1725.

21. Wang, S., Peng, J., and Xu, J. (2011) Alignment of distantly related protein structures: algorithm, bound and implications to homology modeling. Bioinformatics, 27(18), 2537–2545.

22. Ritchie, D. W. (2016) Calculating and scoring high quality multiple flexible protein structure alignments. Bioinformatics, 32(17), 2650–2658.

23. Gilchrist, C. L., Mirdita, M., and Steinegger, M. (2024) Multiple Protein Structure Alignment at Scale with FoldMason. bioRxiv, pp. 2024–08.

24. Needleman, S. B. and Wunsch, C. D. (1970) A general method applicable to the search for similarities in the amino acid sequence of two proteins. J. Mol. Biol., 48(3), 443–453.

25. Altschul, S. F., Madden, T. L., Schäffer, A. A., Zhang, J., Zhang, Z., Miller, W., and Lipman, D. J. (1997) Gapped BLAST and PSI-BLAST: a new generation of protein database search programs. Nucleic Acids Res., 25(17), 3389–3402.

26. Balaji, S. and Srinivasan, N. (2007) Comparison of sequence-based and structure-based phylogenetic trees of homologous proteins: Inferences on protein evolution. J. Biosci., 32, 83–96.

27. Cheng, H., Kim, B.-H., and Grishin, N. V. (2008) MALIDUP: a database of manually constructed structure alignments for duplicated domain pairs. Proteins: Struct., Funct., Bioinf., 70(4), 1162–1166.

28. Cheng, H., Kim, B.-H., and Grishin, N. V. (2007) MALISAM: a database of structurally analogous motifs in proteins. Nucleic Acids Res., 36(Suppl 1), D211–D217.

29. Chandonia, J.-M., Fox, N. K., and Brenner, S. E. (2019) SCOPe: classification of large macromolecular structures in the structural classification of proteins—extended database. Nucleic Acids Res., 47(D1), D475–D481.

30. Boeckmann, B., Dylus, D., Moretti, S., Altenhoff, A., Train, C.-M., Kriventseva, E., Bougueleret, L., Xenarios, I., Privman, E., Gabaldon, T., and Dessimoz, C. (2017) Taxon sampling unequally affects individual nodes in a phylogenetic tree: consequences for model gene tree construction in SwissTree. bioRxiv,.

31. Zhou, N., Jiang, Y., Bergquist, T. R., Lee, A. J., Kacsoh, B. Z., Crocker, A. W., Lewis, K. A., Georghiou, G., Nguyen, H. N., Hamid, M. N., et al. (2019) The CAFA challenge reports improved protein function prediction and new functional annotations for hundreds of genes through experimental screens. Genome Biol., 20, 1–23.

32. Aleksander, S. A., Balhoff, J., Carbon, S., Cherry, J. M., Drabkin, H. J., Ebert, D., Feuermann, M., Gaudet, P., Harris, N. L., et al. (2023) The gene ontology knowledgebase in 2023. Genetics, 224(1), iyad031.

33. Oliveira, G. B., Pedrini, H., and Dias, Z. (2023) TEMPROT: protein function annotation using transformers embeddings and homology search. BMC Bioinf., 24(1), 242.

34. Mensch, A. and Blondel, M. (2018) Differentiable dynamic programming for structured prediction and attention. In Int. Conf. Mach. Learn. PMLR pp. 3462–3471.

35. Van Den Oord, A., Vinyals, O., et al. (2017) Neural discrete representation learning. Adv. Neural Inf. Process. Syst., 30.

36. Zhang, Y. and Skolnick, J. (2004) Scoring function for automated assessment of protein structure template quality. Proteins Struct. Funct. Bioinf., 57(4), 702–710.

37. Murzin, A. G., Brenner, S. E., Hubbard, T., and Chothia, C. (1995) SCOP: a structural classification of proteins database for the investigation of sequences and structures. J. Mol. Biol., 247(4), 536–540.

38. Lefort, V., Desper, R., and Gascuel, O. (2015) FastME 2.0: a comprehensive, accurate, and fast distance-based phylogeny inference program. Mol. Biol. Evol., 32(10), 2798–2800.

39. Tria, F., Landan, G., and Dagan, T. Phylogenetic rooting using minimal ancestor deviation. Nat Ecol Evol 1:0193. (2017).

40. Price, M. N., Dehal, P. S., and Arkin, A. P. (2010) FastTree 2– approximately maximum-likelihood trees for large alignments. PLoS One, 5(3), e9490.

41. Robinson, D. F. and Foulds, L. R. (1981) Comparison of phylogenetic trees. Math. Biosci., 53(1-2), 131–147.

42. Tan, G., Gil, M., Löytynoja, A. P., Goldman, N., and Dessimoz, C. (2015) Simple chained guide trees give poorer multiple sequence alignments than inferred trees in simulation and phylogenetic benchmarks. Proc. Natl. Acad. Sci. U.S.A., 112(2), E99–E100.

43. Consortium, T. U. (2023) UniProt: the universal protein knowledgebase in 2023. Nucleic Acids Res., 51(D1), D523– D531.

44. Brown, P., Pullan, W., Yang, Y., and Zhou, Y. (2016) Fast and accurate non-sequential protein structure alignment using a new asymmetric linear sum assignment heuristic. Bioinformatics, 32(3), 370–377.

45. Singh, D. and Yi, S. V. (2021) On the origin and evolution of SARS-CoV-2. Exp. Mol. Med., 53(4), 537–547.

46. Wen, Z., He, J., and Huang, S.-Y. (2020) Topology-independent and global protein structure alignment through an FFT-based algorithm. Bioinformatics, 36(2), 478–486.

47. Godzik, A. (1996) The structural alignment between two proteins: is there a unique answer?. Protein Sci., 5(7), 1325–1338.

