## Supplementary for "A comprehensive benchmarking study of protein structure alignment tools based on downstream task performance"

#### 1. Alignment quality evaluation on the Malidup dataset

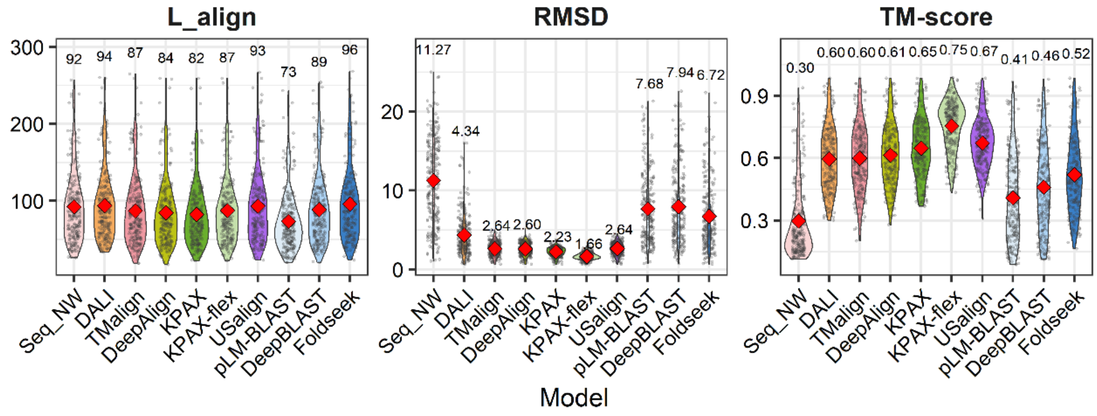


Figure S1. Reference-independent metrics on Malidup dataset.

#### 2. Homology detection statistics at family and fold levels

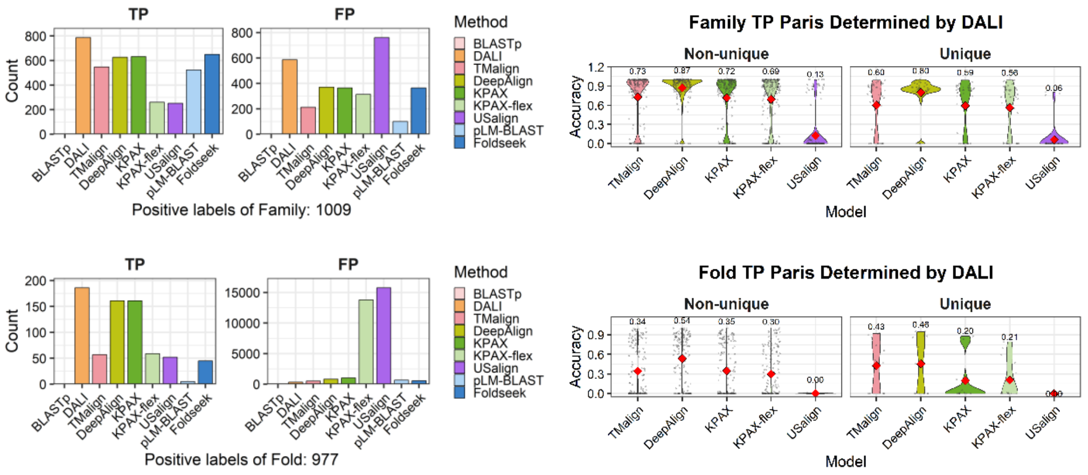


Figure S2. Left: Counts of true positive and false positive pairs determined by different tools on family and fold levels of SCOP140. Right: Alignment accuracy of different tools with respect to the DALI pattern of DALI-unique and -non-unique true possible pairs on family and fold levels.

#### 3. Statistics of proteins in families in SwissTree

| Tree | Family | No. Prot. | Avg. pLDDT | Δ Length (aa.) | Δ Sequence Identity (%) | Δ Structure Identity (%) | Δ RMSD | Δ TM-score |
|---|---|---|---|---|---|---|---|---|
| ST001 | POPDC | 45 | 0.75 | 223-378 (337) | 26-99 | 32-99 | 1.78-5.82 | 0.36-0.88 |
| ST002 | NADPH | 50 | 0.85 | 490-811 (574) | 22-97 | 66-100 | 0.23-3.83 | 0.65-0.99 |
| ST003 | V-ATPase B | 47 | 0.87 | 231-611 (497) | 26-100 | 55-100 | 0.36-2.25 | 0.55-1.0 |
| ST004 | SERINC | 103 | 0.76 | 280-530 (448) | 18-100 | 28-100 | 0.74-5.43 | 0.26-0.99 |
| ST005 | SUMF | 25 | 0.87 | 278-412 (334) | 23-99 | 70-98 | 0.57-2.98 | 0.66-0.99 |
| ST006 | HOX | 95 | 0.62 | 57-493 (271) | 21-100 | 14-98 | 0.24-6.79 | 0.14-0.99 |
| ST007 | RPS | 56 | 0.85 | 99-509 (128) | 16-99 | 27-100 | 0.08-5.03 | 0.32-0.99 |
| ST008 | BAMBI | 35 | 0.61 | 217-664 (281) | 22-100 | 22-100 | 1.67-7.44 | 0.19-0.94 |
| ST009 | Asterix | 34 | 0.60 | 82-198 (111) | 17-100 | 36-100 | 0.79-5.21 | 0.30-0.96 |
| ST010 | CITED | 30 | 0.56 | 103-556 (227) | 24-99 | 14-92 | 2.39-6.95 | 0.14-0.66 |
| ST011 | B-amylase | 131 | 0.83 | 290-810 (548) | 22-99 | 14-100 | 0.04-7.0 | 0.16-1.0 |

Table S1. Description and statistics of proteins included in this study. Protein families with averaged pLDDT less than 0.70 are marked with red color.

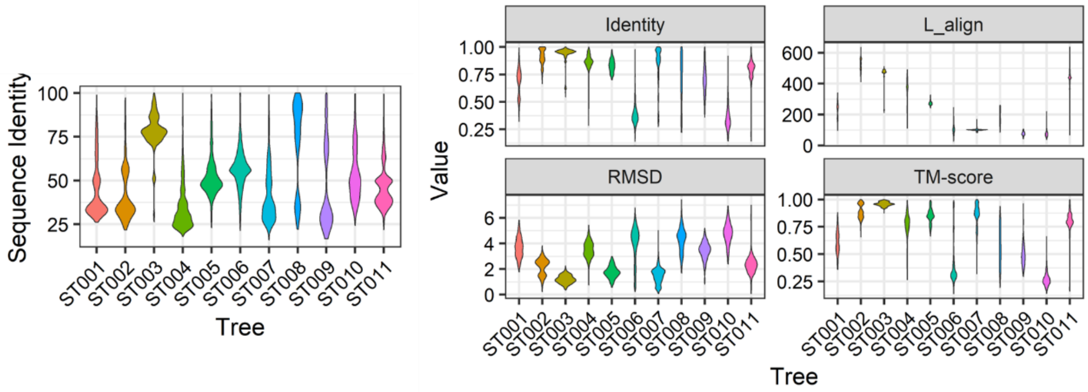


Figure S3. Distribution of sequence identity, SO_identity, L_align, RMSD, and TM-score for each protein families in SwissTree ST001-ST010. Identity on the top-left of the right panel is the structure-based SO_identity, defined as the percentage of aligned regions.

#### 4. Factors that affect ground-truth TCS scores

| Tree | #. Non-binary nodes | #. Paralogs | Size of tree |
|---|---|---|---|
| ST001 | 11 | 26 | 45 |
| ST002 | 9 | 27 | 50 |
| ST003 | 21 | 11 | 47 |
| ST004 | 0 | 57 | 103 |
| ST005 | 0 | 5 | 25 |
| ST006 | 8 | 81 | 95 |
| ST007 | 15 | 11 | 56 |
| ST008 | 0 | 1 | 35 |
| ST009 | 0 | 0 | 34 |
| ST010 | 0 | 12 | 30 |
| ST011 | 0 | 111 | 131 |

Table S2. Numbers of non-binary nodes and paralogs in each protein family. Existence of non-binary nodes and percentage of paralogs higher than 50% are marked with red color.

#### 5. Numbers of consistent columns

| Tree | Δ Length (mean) | Reference | ClustalO | mTMalign | 3DCOMB | KPAX-multi | KPAX-multi-flex | Foldmason |
|---|---|---|---|---|---|---|---|---|
| ST001 | 223-378 (337) | H9G7Q3 | 128 | 18 | 85 | 31 | 119 | 123 |
| ST002 | 490-811 (574) | Q505H9 | 236 | 211 | 241 | 219 | 222 | 201 |
| ST003 | 231-611 (497) | Q4D1S0 | 198 | 192 | 205 | 198 | 200 | 200 |
| ST004 | 280-530 (448) | H2UG27 | 105 | 27 | 35 | 15 | 83 | 85 |
| ST005 | 278-412 (334) | A9V4V4 | 217 | 201 | 216 | 211 | 213 | 196 |
| ST006 | 57-493 (271) | Q1KKV2 | 0 | 0 | OOM | 0 | 0 | 0 |
| ST007 | 99-509 (128) | P54029 | 81 | 70 | 74 | 69 | 67 | 62 |
| ST008 | 217-664 (281) | Q6F2E0 | 114 | 2 | 48 | 12 | 71 | 75 |
| ST009 | 82-198 (111) | Q86H65 | 57 | 9 | 36 | 14 | 28 | 33 |
| ST010 | 103-556 (227) | A0A2J8IVZ1 | 28 | 0 | 23 | 0 | 6 | 19 |
| ST011 | 290-810 (548) | F2DM35 | 0 | 0 | OOM | 4 | 4 | 0 |

Table S3. Numbers of consistent columns determined by different MSA tools.

#### 6. TCS scores in each tree

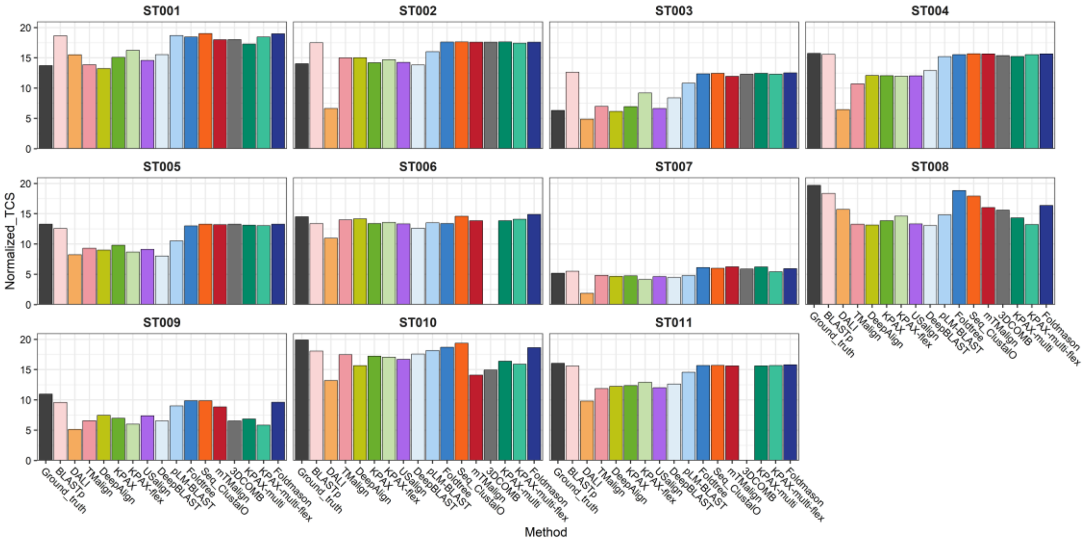


Figure S4. TCS scores of different alignment tools in each tree. Notice that in ST001, ST002, ST003, ST006, and ST007 TCS scores calculated for ground-truth species trees are not the highest because of the existence of non-binary internal nodes.

#### 7. Existence of non-binary internal nodes and paralogs

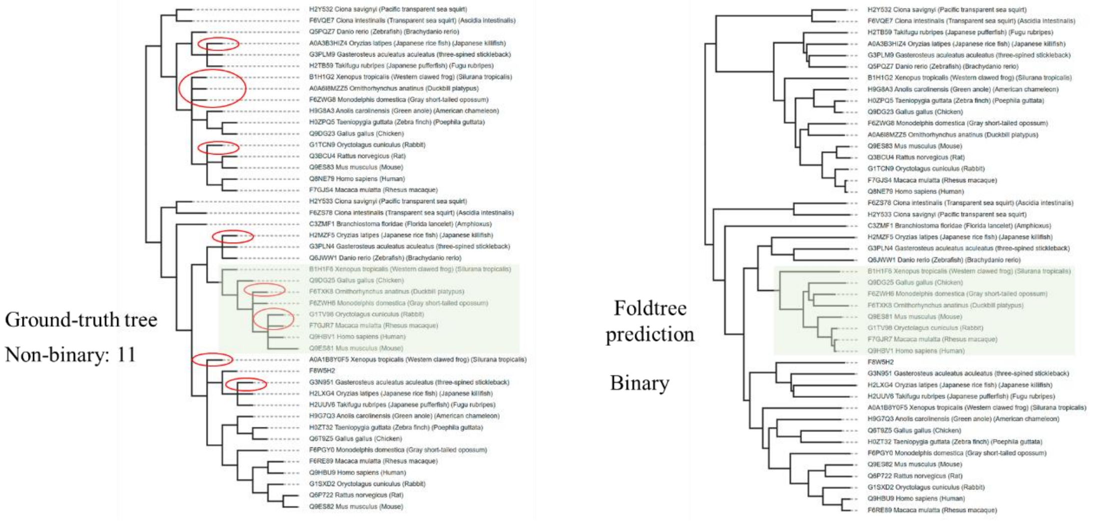


Figure S5. Left: How the number of non-binary internal nodes are determined for a ground-truth species tree (ST001) provide by the SwissTree database. Red circles denote leaves descended from non-binary parents. Right: Similar evolutionary hierarchy is detected by Foldtree in the green box session but with binary structure.

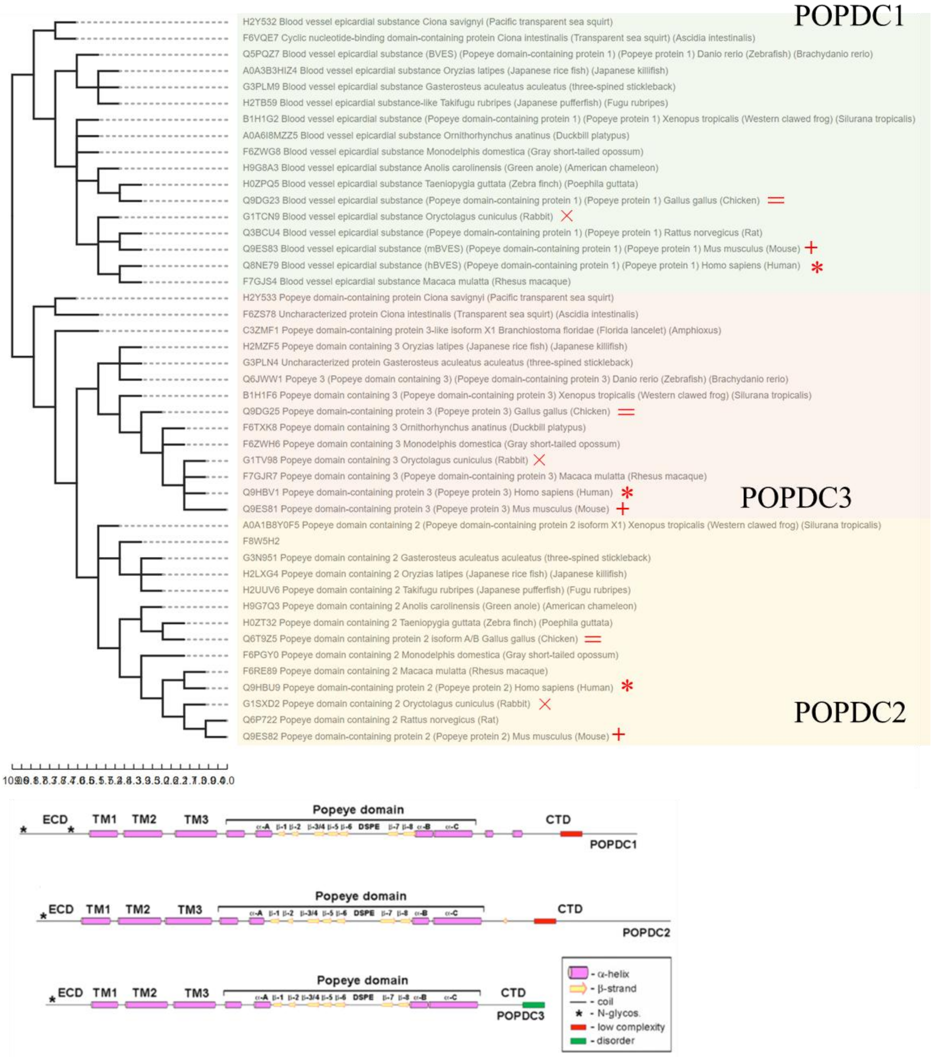


Figure S6. Top: Paralogs exist in protein families. Three different subgroups of the popeye family are boxed with three colors. Each subgroup has proteins belonging to chicken (=), mouse (+), rabbit (×), and human (*). Paralogs are usually generated through duplication of the same protein in one species. Bottom: Cited from https://doi.org/10.1007/s10974-019-09523-z showing the domain differences across the three subgroups.

#### 8. Visualization of multiple alignment of different protein families

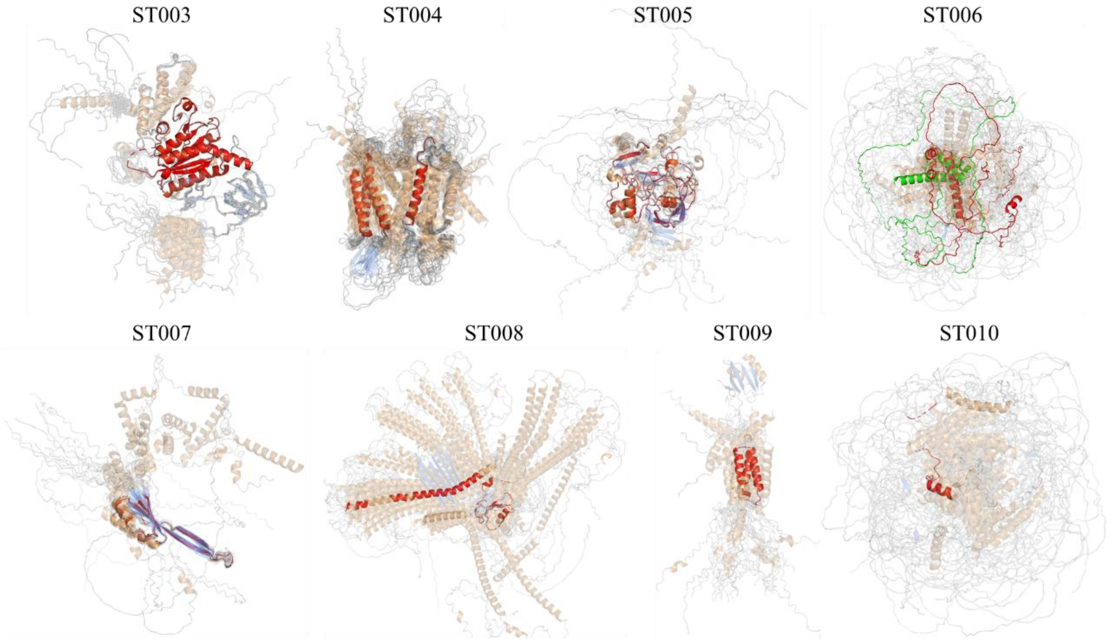


Figure S7. Structure multiple alignment visualization by mTMalign for protein families in SwissTree. Regions include consistent columns determined by Clustal Omega are marked with red color except for ST006, which fails to generate any consensus pattern. For ST006, two proteins are highlighted in green and red, showing that they are not superimposed correctly by mTMalign.

#### 9. Effects of scoring function on function inference of different tools

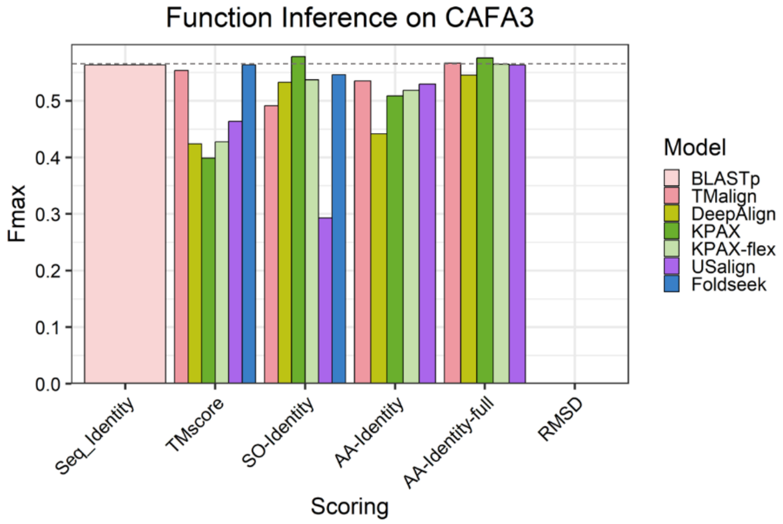


Figure S8. Fmax scores resulting from utilizing different scoring functions for different tools.

#### 10. Correlation between sequence similarity and structure similarity

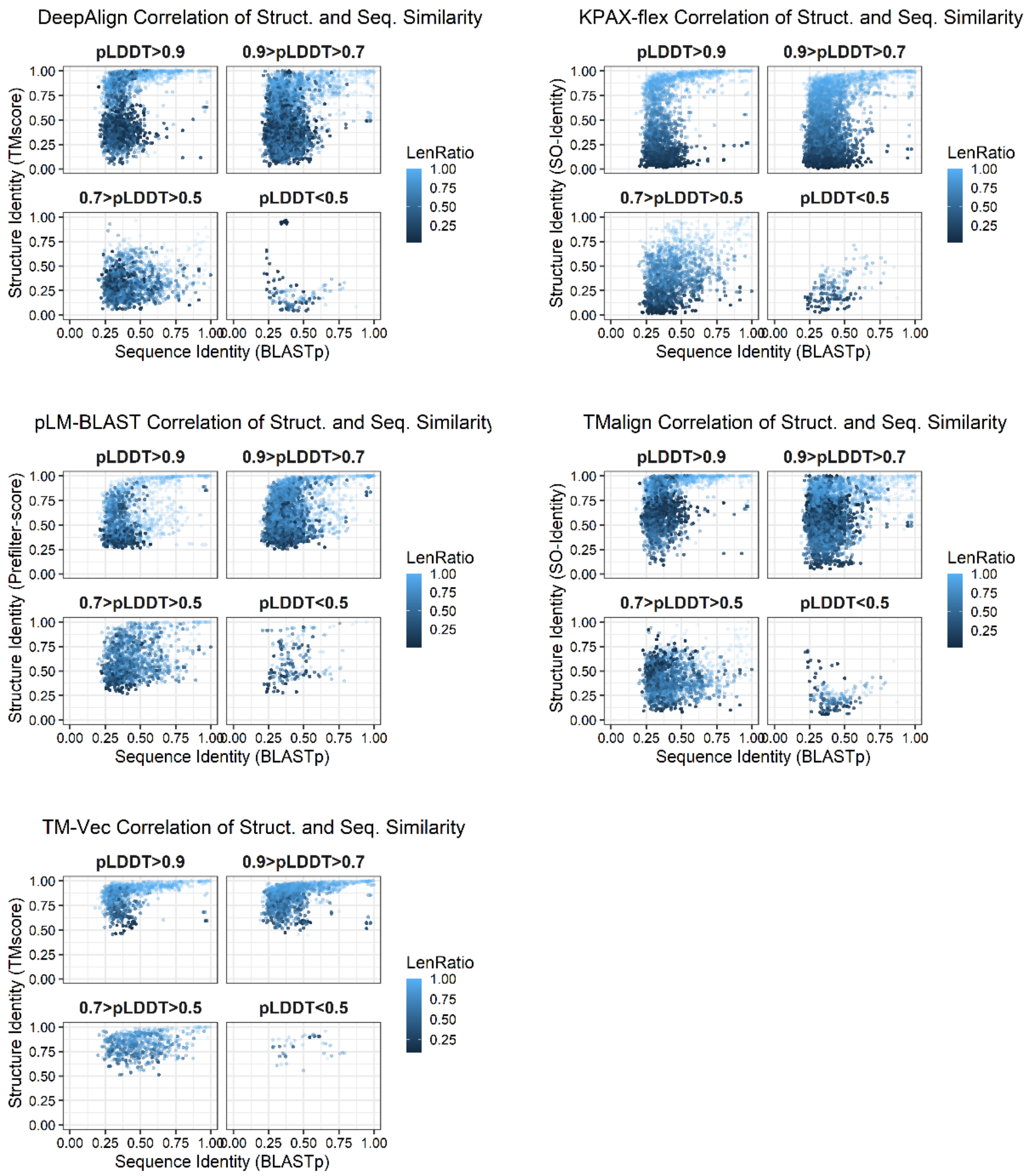


Figure S9. Scatter plots of sequence similarity versus structure similarity of different tools.

##### 11. Runtime and memory consumption of different tools on CAFA3-MF

One query against whole DB [all queries]    Extra time for Indexing and model loading

| Model | QueryLen | Runtime (s) | Memory |
|---|---|---|---|
| BLASTp | <100 (49) | 0.044 [2.2] | 57 MB |
| | 100-300 (417) | 0.132 [55] | 59.5 MB |
| | 300-1000 (570) | 0.405 [231] | 59.8 MB |
| | >1000 (101) | 1.465 [148] | 59.7 MB |
| Diamond | <100 (49) | 0.012 [0.6] | 150.2 MB |
| | 100-300 (417) | 0.001 [0.6] | 150.5 MB |
| | 300-1000 (570) | 0.001 [0.7] | 161.7 MB |
| | >1000 (101) | 0.008 [0.8] | 152.1 MB |
| TMalign | <100 (49) | 0.071 | 2.70 MB |
| | 100-300 (417) | 0.252 | 3.74 MB |
| | 300-1000 (570) | 1.019 | 6.58 MB |
| | >1000 (101) | 5.639 | 14.18 MB |

| Model | QueryLen | Runtime (s) | Memory |
|---|---|---|---|
| TM-Vec | <100 (49) | 0.599 [29] | 16.3 GB |
| | 100-300 (417) | 0.091 [38] | 16.3 GB |
| | 300-1000 (570) | 0.12 [67] | 16.3 GB |
| | >1000 (101) | None | OOM |
| Foldseek | <100 (49) | 0.058 [2.8] | 1.18 GB |
| | 100-300 (417) | 0.024 [9.9] | 2.55 GB |
| | 300-1000 (570) | 0.029 [17] | 2.64 GB |
| | >1000 (101) | 0.210 [21] | 3.20 GB |
| DeepAlign | <100 (49) | 0.491 | 11.5 MB |
| | 100-300 (417) | 1.66 | 15.4 MB |
| | 300-1000 (570) | 6.21 | 26.9 MB |
| | >1000 (101) | 31.8 | 52.3 MB |

| Model | QueryLen | Runtime (s) | Memory |
|---|---|---|---|
| GTalign | <100 (49) | 3.4 [166] | 630 MB |
| | 100-300 (417) | 3.7 [1547] | 616 MB |
| | 300-1000 (570) | 4.4 [2500] | 607 MB |
| | >1000 (101) | 10.3 [1040] | 609 MB |
| DALIX-GPU | <100 (49) | 0.80 | 771 MB |
| | 100-300 (417) | 2.26 | 6.7 GB |
| | 300-1000 (570) | Not Available | OOM |
| | >1000 (101) | | OOM |
| pLM-BLAST | <100 (49) | 0.81 [40] | 10.5 GB |
| | 100-300 (417) | 0.54 [227] | 17.6 GB |
| | 300-1000 (570) | 0.63 [357] | 19.0 GB |
| | >1000 (101) | 0.95 [96] | 17.5 GB |

| Model | QueryLen | Runtime (s) | Memory |
|---|---|---|---|
| Kpax | <100 (49) | 0.133 | 150 MB |
| | 100-300 (417) | 0.299 | 161 MB |
| | 300-1000 (570) | 0.709 | 201 MB |
| | >1000 (101) | 2.37 | 956 MB |
| Kpax-flex | <100 (49) | 0.141 | 150 MB |
| | 100-300 (417) | 0.327 | 161 MB |
| | 300-1000 (570) | 0.821 | 201 MB |
| | >1000 (101) | 2.79 | 956 MB |
| USalign | <100 (49) | 0.170 | 72 MB |
| | 100-300 (417) | 0.462 | 83 MB |
| | 300-1000 (570) | 1.79 | 128 MB |
| | >1000 (101) | 8.78 | 277 MB |

Table S4. Runtime and memory consumption of different tools on CAFA3-MF. Runtime in red denotes that it is estimated for one query protein against the entire target database, and the runtime in square brackets denotes the total query time for all protein in the corresponding length category. Runtime underlined indicates that it requires extra time for indexing and model loading, and such extra time is considered in query time during estimation.

### Implementation and commands

**Needleman-Wunsch dynamic programming**

We provide a custom script for the Needleman-Wunsch algorithm with 1 as the score for a match and -1 for a mismatch and gap penalty.

**BLASTp**

BLAST is downloaded from NCBI (https://ftp.ncbi.nlm.nih.gov/blast/executables/blast+/LATEST/).

Database construction:

makeblastdb -in <TARGETFASTA> -dbtype prot -out <DBNAME>
BLASTp search:
blastp -db <DBNAME> -query <QUERYFASTA> -out result.txt -outfmt 6 -max_target_seqs 1

**TM-align**
TMalign is downloaded from Zhanglab (https://zhanggroup.org/TM-align/TMalign.cpp) and compiled locally. We run TMalign for pairwise comparison with default settings:
TMalign <QUERY.pdb> <TARGET.pdb>

**DALI**
Both the Malidup/Malisam and SCOP140 datasets have included DALI alignment results provided by the authors of these datasets, and those results are generated by the non-sequential version of DALI which is not yet open-source. The public sequential version DaliLite v.5 is downloaded from the DALI server webpage (http://ekhidna2.biocenter.helsinki.fi/dali/README.v5.html), and we use this version for generating alignment patterns for the homology detection task. For the phylogeny reconstruction task, we submit PDB files to the server to obtain multiple alignment results.

**DeepAlign**
DeepAlign is downloaded and compiled from DeepAlign GitHub (https://github.com/realbigws/DeepAlign). We perform pairwise alignment with the following command:
DeepAlign <QUERY.pdb> <TARGET.pdb>

**KPAX**
KPAX is downloaded from the KPAX website (https://kpax.loria.fr). We run pairwise alignment with the flexible setting as an option:
kpax [-flex] <QUERY.pdb> <TARGET.pdb>
For multiple alignment with the flexible setting as an option, we perform:
kpax -multi [-flex] -nopivot <PDB_list>
FastTree <KPAX_MULTI_FASTA> > kpax_multi_tree.nhx

**US-align2**
USalign is downloaded from USalign GitHub (https://github.com/pylelab/USalign). We use batch search and fast mode for the function inference task and use pairwise alignment for other tasks.
Fully non-sequential pairwise alignment:
USalign <QUERY.pdb> <TARGET.pdb> -outfmt 2 -mm 5 -fast
Batch search:
USalign <QUERY.pdb> -dir2 <TARGET_DIR> <TARGET.list> -suffix .pdb -outfmt 2 -mm 5 -fast

**DeepBLAST and TM-Vec**
The DeepBLAST model and checkpoints are downloaded from DeepBLAST GitHub (https://github.com/flatironinstitute/deepblast). Pairwise alignment is performed using a custom

script modified from the DeepBLAST workflow with the model.align(seq1,seq2) function. For database search using TM-vec:
Database build:
tmvec-build-database --input-fasta <TARGET.fasta> --tm-vec-model tm_vec_cath_model.ckpt --tm-vec-config-path tm_vec_cath_model_params.json --device 'gpu' --output target_tmvec_database
TM-Vec search:
tmvec-search --query <QUERY.fasta> --tm-vec-model tm_vec_cath_model.ckpt --tm-vec-config tm_vec_cath_model_params.json --database target_tmvec_database/db --metadata target_tmvec_database/meta.npy --database-fasta <TARGET.fasta> --device gpu --output-format tabular --output TM-Vec_tabular.txt --output-embeddings tmvec_query_embeddings.npy

**pLM-BLAST**
The pLM-BLAST model is downloaded from pLM-BLAST GitHub (https://github.com/labstructbioinf/pLM-BLAST). Pairwise alignment is performed using a custom script modified from the pLM-BLAST workflow with
the embedding_to_span(emb1,emb2) function.
Database construction:
python embeddings.py start <QUERY.fasta> query_plmblast_database -embedder pt --gpu -bs 0 --asdir
python embeddings.py start <TARGET.fasta> target_plmblast_database -embedder pt --gpu -bs 0 --asdir
Database pre-filtering search:
python scripts/plmblast.py target_plmblast_database query_plmblast_database > all.oc.json -oc

**Foldseek**
The Foldseek model and pipeline are downloaded from Foldseek GitHub (https://github.com/steineggerlab/foldseek). Foldseek database search is performed with:
foldseek easy-search <QUERYDB> <TARGETDB> allvsall.tsv tmp --format-output 'query,target,fident,alntmscore' -a -e inf --exhaustive-search

**Foldtree**
The Fold_tree model and workflow are downloaded from Foldtree GitHub (https://github.com/DessimozLab/fold_tree).
snakemake --cores 4 --use-conda -s ./workflow/fold_tree --config folder=./myfam filter=False custom_structs=True

**Clustal Omega**
Clustal Omega version 1.2.4 is used. Commands for multiple alignment and tree construction are as follows:
clustalo -i <FASTA> -o aln.fasta -v
FastTree aln.fa > clustalo_tree.nhx

**mTMalign**

mTMalign is downloaded from the URL in the original paper (http://yanglab.nankai.edu.cn/mTM-align). The following are commands for multiple alignment and tree construction:

mTMalign -i <PDBLIST>
sed -i 's/.pdb//g' result.fasta
FastTree result.fasta > mTMalign_Tree.nhx

**3DCOMB**

The 3DCOMB program is included in the DeepAlign tool. The following are commands for multiple alignment and tree construction:

3DCOMB -i <QUERY_LIST> -o aln
sed -i 's/.pdb//g' aln.ali
FastTree aln.ali > deepalign3DCOMB_tree.nhx

**Foldmason**

The Foldmason model and pipeline are downloaded from Foldmason GitHub (https://github.com/steineggerlab/foldmason). Multiple alignment and tree construction are conducted with the following commands:

foldmason createdb <PDBFOLDER>/* foldmason
foldmson structuremsa foldmason result
FastTree result_aa.fa > foldmason_tree.nhx
